# ERG1A K^+^ Channel Increases Intracellular Calcium Concentration through Modulation of Calsequestrin1 in C_2_C_12_ Myotubes

**DOI:** 10.1101/2023.12.04.569937

**Authors:** Gregory H. Hockerman, Evan Pratt, Shalini Guha, Emily LaVigne, Clayton Whitmore, Omar Khader, Natalie McClure, Sandra Zampieri, Jennifer Koran, W-H Wang, Amber L. Pond

## Abstract

The ERG1A K^+^ channel modulates the protein degradation that contributes to skeletal muscle atrophy by increasing intracellular calcium concentration ([Ca^2+^]i) and enhancing calpain activity, but the mechanism by which the channel regulates the [Ca^2+^]i is not known. Here, we have investigated the effect of human ERG1A (HERG) on [Ca^2+^]i in C_2_C_12_ myotubes, using Fura-2 calcium assays, immunoblot, RT-qPCR, and electrophysiology. We hypothesized that HERG would modulate L-type calcium channel activity, specifically the Cav1.1 channel known to carry signal from the sarcoplasmic membrane of skeletal muscle to the sarcomeres of the myofibrils. However, we find that HERG has no effect on the amplitude of L-type channel current nor does it affect the mRNA levels nor protein abundance of the Cav1.1 channel. Instead we find that, although the rise in [Ca^2+^]i (induced by depolarization) is greater in myotubes over-expressing HERG relative to controls, the difference between the KCl-stimulated Ca^2+^ increase in control and HERG over-expressing cells cannot be accounted for by L-type channel mediated Ca^2+^ influx, which suggests that HERG could modulate excitation coupled calcium entry (ECCE). Indeed, the HERG-enhanced increase in [Ca^2+^]i induced by depolarization is blocked by 2-APB, an inhibitor of ECCE (and SOCE). Further, we show data suggesting that HERG also modulates the activity of ryanodine receptors, a component of ECCE, as well as store operated calcium entry (SOCE). Therefore, we investigated the effect of HERG on calsequestrin1, a calcium buffering/binding protein known to modulate ryanodine receptor 1 and store operated Ca^2+^ entry activities. Indeed, we find that calsequestrin1 mRNA levels are decreased 0.83-fold (p<0.05) and the total protein abundance is lowered 77% (p<0.05) in myotubes over-expressing HERG relative to controls. In summary, the data show that ERG1A overexpression modulates [Ca^2+^]i in skeletal muscle cells by lowering the abundance of the calcium buffering/binding protein calsequestrin1.

## INTRODUCTION

The *ether-a-go-go related gene* (*erg1*) encodes a number of ERG1 K^+^ channel alpha subunit alternative splice variants. Two ERG1 splice variants, which have been cloned from both human ERG1 (HERG1A and 1B; London et al., 1997) and mouse *Erg1* (*Merg1a* and *1b*; Lees-Miller et al., 1997) cDNA libraries, form a heteromultimeric channel in mammalian heart. This ERG1A/1B channel has been shown to produce I_Kr,_ a current which is partially responsible for late phase repolarization of the cardiac action potential (Jones, et al., 2004; Curran et al., 1995). We have shown that the ERG1A K^+^ channel subunit contributes to skeletal muscle loss in atrophic situations through up-regulation of ubiquitin proteasome proteolysis (UPP); we do not detect ERG1B in skeletal muscle (Wang et al., 2006). Specifically, our labs have reported that the ERG1A K^+^ channel is up-regulated in skeletal muscle of mice undergoing atrophy as a result of hind limb unweighting (Wang et al., 2006), tumor-induced cachexia (Wang et al., 2006; Zampieri et al., 2021), and denervation (Anderson et al., 2021). Moreover, we have reported that: 1) pharmacological and genetic block of ERG1A function inhibit skeletal muscle atrophy induced by hind limb suspension (Wang et al., 2006); 2) atrophy is induced in mouse muscle by ectopic expression of wild type *Erg1a* (Wang et al., 2006); and 3) abundance of the UPP E3 ubiquitin ligase MuRF1 (but not ATROGIN1) and overall UPP activity are increased in mouse skeletal muscle by ectopic overexpression of *Erg1a* (Hockerman et al., 2014; Pond et al., 2013; Wang et al., 2006). Further, we have shown that the ERG1A K^+^ channel protein is expressed at low abundance in C_2_C_12_ myotubes and that augmentation of its expression in these cells, as in mouse skeletal muscle, will induce a decrease in cell size (i.e., myotube area) and an increase in the abundance of the E3 ubiquitin ligase MuRF1, but not the E3 ligase ATROGIN1 (Whitmore et al., 2020). Finally, human Erg1A (HERG) overexpression in C_2_C_12_ myotubes induces a significant increase in basal intracellular calcium ([Ca^2+^]i) and calpain activity at 48 hours after transduction (Whitmore et al., 2020). Thus, upregulation of ERG1A contributes to skeletal muscle atrophy *in vivo*, and ectopic expression of HERG in C_2_C_12_ myotubes induces an atrophic phenotype *in vitro*. Given the potentially important role of Ca^2+^ in modulation of skeletal muscle health and atrophy, we sought to understand the mechanism whereby HERG expression elevates basal [Ca^2+^]i in C_2_C_12_ myotubes (Whitmore et al., 2020).

Intracellular Ca^2+^ is essential for excitation-contraction coupling (ECC) in skeletal muscle. Specifically, Cav1.1 L-type voltage-gated Ca^2+^ channels (also known as dihydropyridine receptors, DHPR) are activated by the depolarizing action potentials that spread longitudinally along the muscle sarcolemma and inwardly into the myofibers along the t-tubules, where they are juxtaposed to ryanodine receptors 1 (RyR1). Through a physical interaction, the RyR1 and Cav1.1 together allow release of Ca^2+^ from the sarcoplasmic reticulum (SR) into the cytosol through the RyR1 channel. This increase in [Ca^2+^]i (μM range) triggers skeletal muscle contraction, which is reversed when Ca^2+^ is “pumped” back into the SR by Ca^2+^-ATPase (SERCA) (reviewed in Cho et al., 2017). Smaller increases in [Ca^2+^]i (nM range) also occur in cells and these modulate muscle cell events such as upregulation of muscle thermogenesis in non-contracting muscle in response to cold (Bal et al., 2012) and increase of both fatigue resistance and mitochondrial biogenesis (Bruton et al., 2010). Localized Ca^2+^ concentration fluctuations also serve as second messengers, modulating numerous signaling systems (Tu et al., 2016). Thus, [Ca^2+^]i is affected by mechanisms other than the physical interaction between Cav1.1 and RyR1 that occurs in response to sarcolemmal membrane depolarization (i.e., ECC). One mode of Ca^2+^ modulation which results in relatively small changes in [Ca^2+^]i is termed excitation coupled Ca^2+^ entry (ECCE). This Ca^2+^ influx is activated in response to prolonged and repetitive depolarization (e.g., by treatment with mM KCl *in vitro*) through a Cav1.1 and RyR1-mediated interaction, but it is insensitive to lower concentrations (≤10 μM) of the L-type current inhibiting reagent nifedipine (Bannister et al., 2009; Yang et al., 2007). It is inhibited by 2-APB, which also blocks store operated Ca^2+^ entry (SOCE) and IP_3_ receptors at the concentrations we used (100 μM), but is not affected by Ca^2+^ store depletion (Cherednichenko et al., 2004; Hurne et al., 2005). ECCE requires both RyR1 and Cav1.1 proteins (Cherednichenko et al., 2004) and L-type channel pore permeation appears to contribute to this calcium ion passage (Bannister et al., 2009); however, it is still likely that this extracellular entry mechanism may require another (as yet unidentified) sarcolemmal Ca^2+^ channel or other auxiliary units (Dirksen 2009). To date, it has been shown that Orai1 (Lyfenko and Dirksen 2008) and TRPC3 (Lee et al., 2006) are not candidates for this role. Calcium is also brought into the cell from the extracellular milieu through SOCE. SOCE is inhibited by depolarization and is activated by depletion of intracellular Ca^2+^ stores (i.e., the SR) (Kurebayashi & Ogawa 2001). Specifically, SR depletion results in dimerization of two SR membrane STIM1 proteins which are a major component of SOCE. Once dimerized, the STIM1 proteins then translocate through the SR membrane to interact with and open the sarcolemmal membrane Orai1 Ca^2+^ channel through which the Ca^2+^ moves into the cell from the extracellular milieu (Roos et al., 2005; Feske et al., 2006; Vig et al., 2006).

Here, we show that, relative to controls, the abundance of the Ca^2+^ buffering protein calsequestrin1 (CaSeq1) is significantly lower in HERG over-expressing myotubes. Because RyR1 and SOCE activities are inhibited by CaSeq1 at higher concentrations of [Ca^2+^]i (Jeong et al., 2021; Zhang et al., 2016; Wang et al., 2015; Wei et al., 2009), the down regulation of CaSeq1 in response to HERG overexpression could produce an increase in [Ca^2+^]i by enhancing the open probability of the RyR1 channel, which would increase Ca^2+^ outflow from SR stores. Indeed, we observe that overexpression of HERG increases RyR1 activity. The activation of RyR1 could also potentially stimulate the ECCE, which would increase influx of calcium to the cytosol from the extracellular milieu. We do report here an increase in ECCE activity with HERG overexpression. Additionally, we note that HERG overexpression enhances SOCE activity, which may perhaps occur as a direct effect on a SOCE component or from the decrease in [Ca^2+^]_SR_ resulting from the HERG-related increased RyR1 activity. Indeed, we report here that the activities of RyR1 and the ECCE as well as the SOCE pathway are enhanced in myotubes overexpressing HERG.

## MATERIALS AND METHODS

### Cell Culture

C_2_C_12_ cells (ATCC; Manassas, VA) were cultured in Dubelco’s Minimum Essential Medium (DMEM; ThermoFisher Scientific; Waltham, MA) containing 25 mM glucose, 4 mM L-glutamine and 3.7 g/L sodium bicarbonate and supplemented with 10% fetal bovine serum (FBS; ThermoFisher Scientific) within a 37°C incubator containing 6.5% CO_2_. Differentiation of myoblasts into myotubes was induced by switching to culture medium as described here except that the 10% FBS was replaced with 2% heat-inactivated horse serum (HS). After replacement of medium, cells were allowed to differentiate for 7-8 days at which point myotubes were transduced and allowed to incubate another 48 hours with virus (i.e., another 2 days to produce 9-10 days cultured cells).

### Antibodies

All antibodies (except the antibodies used in Supplemental Data 3, SD3) were diluted in phosphate buffered saline (PBS, pH 7.4) with 5% normal goat serum, 0.1% Triton X-100, and 0.01% sodium azide. For the Calsequestrin1 immunoblot in Figure 8, the Calsequestrin1 D-10 (sc-137080; Santa Cruz, Dallas, TX) mouse monoclonal primary antibody was diluted in this buffer at 1:100. To detect Cav1.1 α1 protein, the DHPR α1 primary antibody (MA3-920; ThermoScientific) was used at a 1:500 dilution. For GAPDH protein immunoblots, we probed the membranes with a monoclonal antibody (Clone 6C5, MilliporeSigma; St. Louis, MO) diluted 1:8000 and the β-tubulin antibody (T8328, Sigma) was diluted 1:2000. The alkaline phosphatase conjugated goat anti-mouse IgG antibody (BioRad; Hercules, CA) was used as secondary antibody (with all of the primary antibodies described above) at 1:10,000 in blotting buffer (0.2% non-fat dry milk or casein and 0.1% Tween-20 in Tris Buffered Saline, pH 7.4; Sigma; St. Louis, MO). For the immunoblot in SD3, the calsequestrin1-specific primary antibody 26665-1-AP (ProteinTech; Rosemont, IL) was diluted 1:1000 in 0.2% non-fat dry milk in Tris-buffered saline with 0.1% tween-20, 5% NGS and 0.1% sodium-azide (pH 7.4, TTBS) followed by washing in TTBS and then incubation in secondary antibody (goat anti-rabbit IgG-Alkaline Phosphatase; 65-6122 Sigma) diluted 1:1000 in 0.2% non-fat dry milk in TTBS.

### Virus

The human ERG1A construct (Lin et al., 2010), was cloned into a viral cassette provided by ViraQuest, Inc. (North Liberty, IA) and then cloned by this company into the VQad CMV adenovirus (containing GFP expression apparati). The appropriate control virus was also purchased from this company and used for the control transfections. Both sets of viral particles were maintained at −80°C in small aliquots until used.

### Viral Transduction of C_2_C_12_ Myotubes

HERG and control viral particles were added to DMEM with 2% horse serum (Gibco/ThermoFisher; Waltham, MA) and mixed. This suspension was used to treat C_2_C_12_ myotubes (cultured as described above) at 200 multiplicity of infection (MOI) as pre-determined by titration, EXCEPT that the myotubes used for the calsequestrin1 and GAPDH immunoblots in Figure 7 were treated with 400 MOI. The virus treated plates were gently agitated to properly disperse the viral particles and then returned to the incubator. Viral expression was monitored by GFP fluorescence and detected at 24 hours post transfection; it appeared to plateau at 48 hours, so this is when procedures were performed.

### Intracellular Calcium Concentration Determinations

A Fura-2 QBT Calcium Kit (Molecular Devices; San Jose, CA) was used to assess [Ca^2+^]i according to manufacturer’s instructions. Briefly, myotubes were grown in the wells of a 96-well plate as described earlier. The myotubes were then loaded with Fura-2 QBT by incubation with 1:1 Gibco Opti-MEM (Life Technologies; Carlsbad, CA) and Fura-2 QBT (dissolved at recommended concentration) in modified Krebs-Ringer Buffer with HEPES (mKRBH; 0.05% fatty acid free bovine serum albumin in 134 mM NaCl, 3.5 mM KCl, 1.2 mM K_2_HPO_4_, 0.5 mM MgSO_4_, 1.5 mM CaCl_2_, 5 mM NaHCO_3_, 10 mM HEPES) for 1 hour at room temperature (RT) in the dark. This medium was then replaced with Fura-2 QBT solution (in mKRBH) containing either a treatment (as described below) or vehicle (as control) and incubated at RT in the dark for 30 minutes. A Synergy Mx Microplate Reader (BioTek; Winooski, VT) was used in conjunction with Genesis 2.0 software to excite the samples at 340 nm and 380 nm and to measure emission at 508 nm.

The 340/380 nm emission ratio was determined for all recorded time points for each reaction (i.e., plate well). The average of the background 340/380 ratios (prior to addition of treatment) was determined for each plate well and that was subtracted from the 340/380 ratio for each time point (after addition of treatment) recorded for that well to normalize data to background as an assessment of [Ca^2+^]i flux. The normalization to background *(recorded prior to addition of treatment)* is necessary to correct for the significant differences in basal [Ca^2+^]i levels between control and HERG-overexpressing myotubes. Indeed, we have recorded/reported significantly higher [Ca^2+^]i levels in HERG overexpressing versus control myotubes at 48 hours post transduction (Whitmore et al., 2020), the time at which we performed our Fura2 assays here to ensure HERG overexpression.

### Depolarization Assays

Myotubes were cultured and treated with either control virus or virus encoding HERG as described earlier. At 48 hours post viral transduction, these myotubes were loaded with Fura-2 QBT and then treated with either vehicle or drug for 30 minutes. The drug was either: nicardipine (10 μM, an L-type Ca^2+^ current inhibitor), nifedipine (10 μM, an L-type Ca^2+^ current inhibitor), thapsigargin (Tg, 1 μM, an inhibitor of SERCA1), 2-aminoethoxydiphenyl borate (2-APB; 100 μM to block SOCE, ECCE, and IP_3_R; Sigma), or astemizole (1 nM, a HERG channel blocker). The fluorescence assay for [Ca^2+^]i (as described above) was monitored for one minute (to determine background) and then the cells were depolarized by addition of 100 mM KCl and assayed for an additional 2.5 minutes (in the continued presence of the KCl).

### Ryanodine Receptor Assays

Myotubes were cultured and treated with either control virus or virus encoding HERG as described earlier. At 48 hours post viral transduction, the myotubes were loaded with Fura-2 QBT and then treated for 30 minutes with ryanodine (100 μM, as determined by a dose-response curve to completely block caffeine induced Ca^2+^ release; data not shown) to block RyR1. The Fura-2 QBT [Ca^2+^]i assay (as described above) was started for one minute and then the cells were treated with caffeine (10 mM) to activate RyR1 and finally assayed for an additional 1.5 minutes.

### SOCE Assays

Myotubes were cultured and treated with either control virus or virus encoding HERG as described earlier. At 48 hours post viral transduction, the myotubes were treated with either vehicle or 2-APB (100 μM; to block SOCE activity) for 30 minutes. The cells were then assayed for [Ca^2+^]i using Fura-2 QBT as described above. After 1 minute of assay, an aliquot of vehicle or Tg (to “empty” the SR stores; final 1μM) was dispensed into the wells and [Ca^2+^]i was assayed for an additional 20 minutes to ensure Ca^2+^ levels were consistently low. At 20 minutes, Ca^2+^ (final 2.5 mM CaCl_2_) was added to activate SOCE and the cells were assayed for an additional 10 minutes. The 340/380 nm ratio measured in each well at each time point for two minutes prior to the addition of calcium (rather than the addition of Tg) was averaged and subtracted from the 340/380 nm ratio for each time point after addition of CaCl_2_ to normalize data to background as an assessment of [Ca^2+^]i flux. Again, this normalization was done to correct for differences (i.e., increases) in basal calcium levels that occur in response to HERG overexpression during the 48 hours prior to assay.

### RNA Isolation and RT-qPCR

Total RNA was extracted from control and HERG-expressing C_2_C_12_ myotubes using Trizol reagent (Life Technologies; Carlsbad, CA) according to manufacturer’s instructions followed by chloroform solubilization and ethanol precipitation.

RT-qPCR (quantitative reverse transcriptase PCR) was then performed using a Luna® Universal One-Step RT-qPCR Kit (New England Biolabs; E3005) per manufacturer’s instructions and primers for the CaSeq1 gene of interest (IDT; San Jose, CA) along with primers for ERG1A (Invitrogen; Waltham, MA) and the GAPDH “housekeeping gene” (IDT) (Table 1). A CFX BioRad OPUS 96 real time PCR system was used to detect SYBR green fluorescence as a measure of amplicon. Changes in gene expression were determined using the Livak method to normalize the gene of interest to the “housekeeping gene.” No template controls (NTC) and no-reverse transcriptase (no-RT) controls were also assayed along with our samples of interest in each set of PCR reactions to assess and ensure assay quality.

**Table 1.**
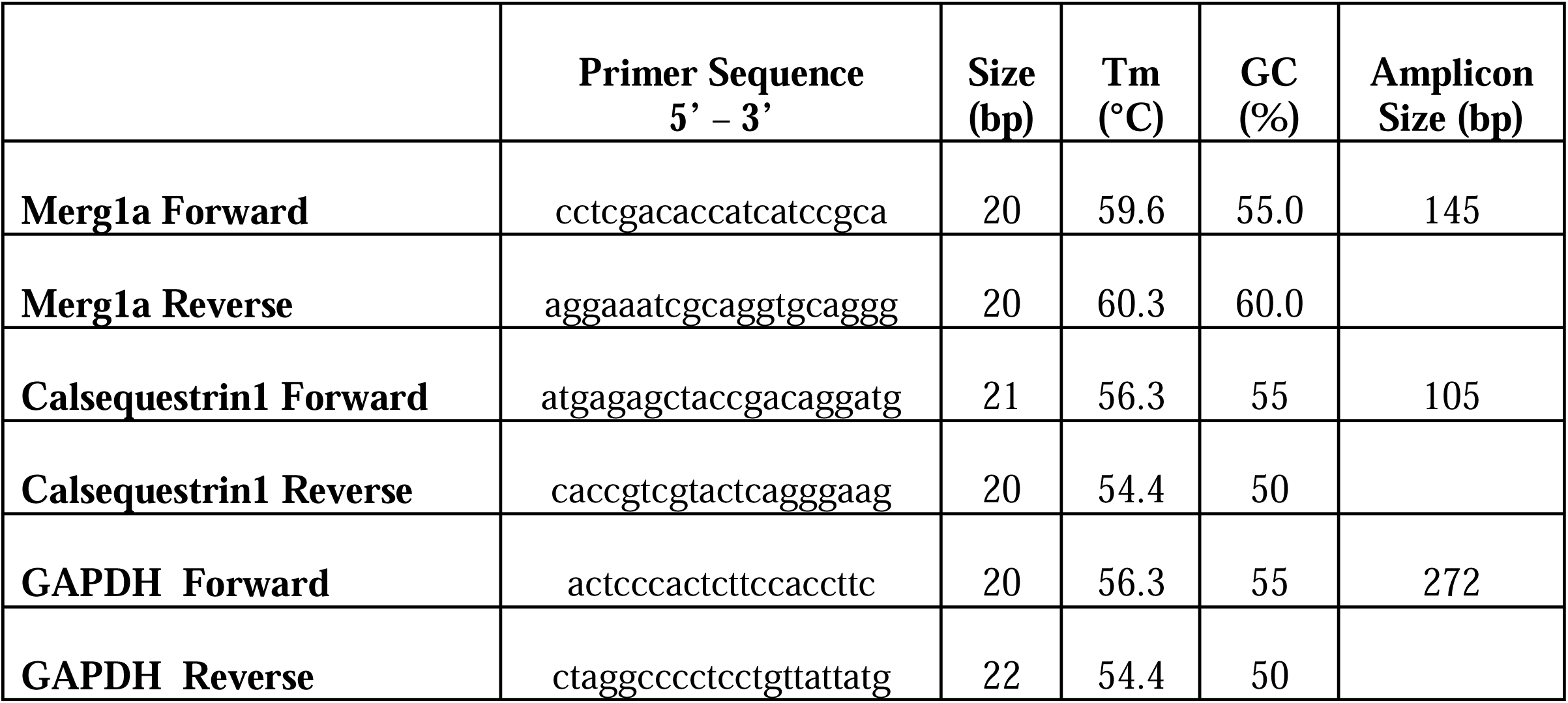
Primer sequences for RT-qPCR.

### Cell Lysate Preparation and Protein Assay

C_2_C_12_ cells were differentiated in 10 cm plates for 7-8 days and then transduced with either control or HERG-encoded virus and incubated an additional 48 hours. Membrane proteins were extracted from myotubes by gently washing tissue culture plates twice with room temperature PBS and then scraping cells from the plates treated with 200 μL of cold Tris-EDTA buffer (10 mM, 1 mM respectively, pH 7.3; Sigma) containing 2% Triton-X100 (Sigma) and protease inhibitors: 0.5 mM pefabloc, 1 mM benzamidine, 1 mM iodoacetamide, 1 mM 1,10-phenanthroline, and a commercial protease cocktail tablet (Pierce A32955 [ThermoFisher] used per manufacturer instructions). Cell suspensions were triturated on ice with a 27.5-gauge needle and 1 mL syringe (∼30 sec) and then again with a 30.5-gauge needle and 1 mL syringe (∼30 sec). The samples were then allowed to sit within ice in a refrigerator for 20 minutes to enhance membrane solubilization. The cell lysates were then triturated with an Eppendorf pipette (200 uL tip) and centrifuged at 12,000xg for 10 minutes to remove solid material. Sample supernatants were collected and stored at −80°C.

### Protein Determination

To determine sample protein concentration, a standard curve was developed using BSA (Sigma) and then a DC Protein Assay kit (BioRad; Carlsbad, CA) was used to develop color in the standards and samples from which protein content was determined by linear regression using Excel (Microsoft 365; Redmond, WA).

### Immunoblot

For immunoblots, aliquots of equal protein content (40 ug, except Fig. SD3 C,E) were boiled for 5 minutes with sample dilution buffer (5X SDB; 0.3 M Tris [pH 6.9], 50% glycerol, 5% SDS, 0.5 M dithiothreitol, and 0.2% bromophenol blue). Samples were then electrophoresed through a 4-20% polyacrylamide (SDS-PAGE) gradient gel, transferred to PVDF membrane (BioRad; Hercules, CA), and immunoblotted using antibodies specific for either: calsequestrin1, Cav1.1 (also called DHPR), GAPDH, or β-tubulin. Secondary antibodies conjugated with alkaline phosphatase were used with an ImmunStar™-Alkaline Phosphatase (AP) Western Chemiluminescent Kit (Bio-Rad) for signal development. To ensure that lanes were loaded equally, after development for CaSeq1 or Cav1.1 signal, PVDF membranes were incubated (30 minutes at room temperature) in stripping buffer (62.5mM Tris Cl [pH 6.9], 2% SDS, 0.695% BME in water), rinsed with 0.1% Tween 20 in 20 mM Tris (TTBS, pH 7.4) and re-probed with GAPDH or β-tubulin antibody (see next section for imaging details). To further confirm protein loading equity, Coomassie Blue stain solution (0.1% Coomassie Blue R-250 in 45% methanol and 10% acetic acid in water) was used to stain the membranes overnight, after which these were destained with 50% methanol and 10% acetic acid in water.

### Immunoblot Imaging

ImageJ (NIH; NC) was used to determine the optical densities of the protein bands: a single region of interest (ROI) which encompassed each protein band was outlined and this ROI was used to demarcate each protein band and a background area above each band. The average uncalibrated optical density (OD) was determined for each ROI and the background OD was subtracted from the OD of each target protein band (e.g., Cav1.1 and CaSeq1). The same was done for the internal reference protein. Then to correct for any differences in sample loading, for each sample a ratio was calculated by dividing the background-corrected OD for each target protein by the background-corrected OD for the “housekeeping protein” for each sample.

### Electrophyisology: Barium Current Densities

C_2_C_12_ myoblasts were seeded in 35 mm cell culture dishes, differentiated to myotubes, and transduced with either control eGFP encoded virus or HERG encoded virus as described. Micropipettes were pulled from borosilicate capillaries to an inside diameter of 3-5 microns using a Sutter P-87 pipette puller, and polished with a Narishige MF 830 microforge. The pipette solution contained (mM): 180 NMDG, 40 HEPES, 4 MgCl_2_, 12 phosphocreatine, 5 BAPTA, 2 Na_2_ATP, 0.5 Na_3_GTP, 0.1 leupeptin, and pH was adjusted to 7.3. The extracellular solution contained (mM): 140 NaCl, 20 CsCl_2_, 10 BaCl_2_, 10 HEPES, 10 glucose, 10 sucrose, 1 MgCl_2_, and pH was adjusted to 7.4. Voltage dependent barium currents were measured under voltage clamp using an Axopatch 200B amplifier (Axon Instruments). Data were sampled at 10 kHz and filtered at 1 kHz. Cells were held at −80 mV and stepped to test voltages ranging from −60 mV to +50 mV in 10 mV increments. Current densities (pA/pF) were calculated by dividing the largest current amplitude for each cell, regardless of test voltage, by the whole cell capacitance.

### Statistics

Net area under the curve (AUC) data were calculated and all statistical analyses were performed using GraphPad Prism 9 (Dotmatics; Boston, MA). A Student’s T-test was used to analyze the optical density data from the immunoblots, the Ca^2+^ current densities, and the HERG and calsequestrin1 gene expression data. A one-way ANOVA was used to analyze the AUC data from the RyR1 and SOCE activity studies as well as the fold changes in Cav1.1 gene expression data. When significant differences were found, means were separated by Tukey’s test. A 2×3 ANOVA was used to compare drug effects on [Ca^2+^]i in response to viral treatment (HERG or control). For the samples depolarized with KCl, the AUCs per time unit were determined and analyzed with a two-way ANOVA design for repeated measures and the interaction between HERG and treatment was examined for statistical significance. The drug sensitive fluorescence ratios (i.e., the mean difference between the activated and activated-blocked samples) were determined by subtracting the mean 340/380 ratio of the reagent activated-blocked sample from the mean 340/380 ratio of the reagent activated sample for each timepoint. The standard error of the mean difference (SEMD) was calculated using equation 1:

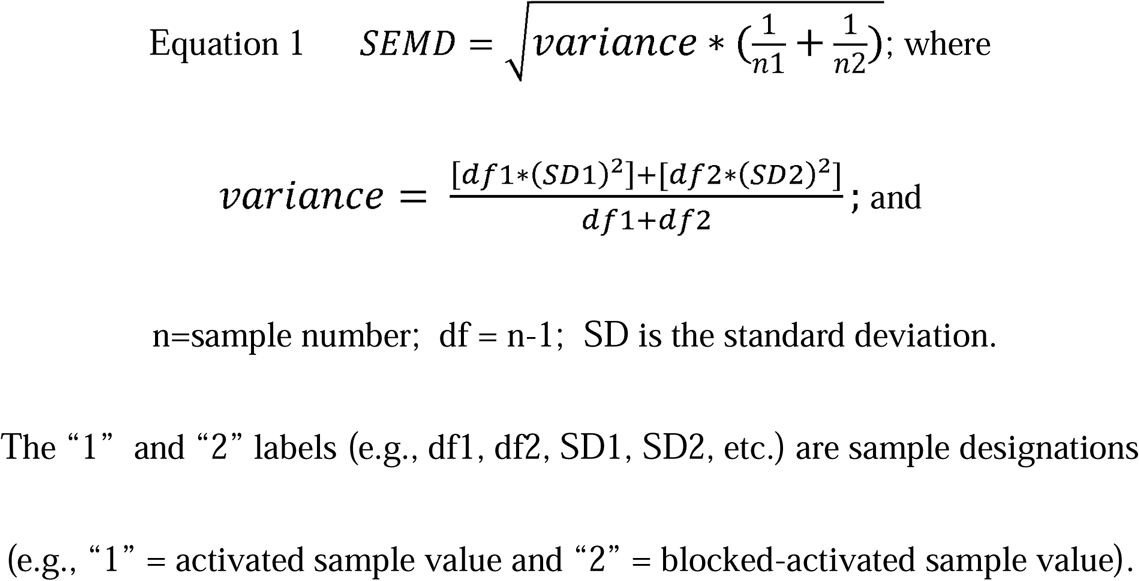

The mean difference and SEMD were used to estimate p values using a comparison of means calculator (MedCalc Software Ltd.; Belgium; https://www.medcalc.org/calc/comparison_of_means.php).

## RESULTS

### The HERG channel affects [Ca^2+^]i by modulation of Excitation Coupled Calcium Entry

#### Depolarization with KCl

HERG-expressing myotubes (Whitmore et al., 2020 validates this viral expression model; also SD1) show a significantly higher intracellular Ca^2+^ concentration than control cells for ∼1.5 minutes after depolarization with 100 mM KCl (Fig. 1A). *Nifedipine Block.* Nifedipine (10 μM to block L-type channel conduction, but not ECCE which is blocked by 50 μM; Bannister et al., 2009) has a strong blocking effect on the increase in [Ca^2+^]i that normally occurs in response to depolarization in the control cells, inhibiting ∼67.5% of the rise in [Ca^2+^]i at 20s and then lowering it substantially (over 90% blocked) from 40s to 120s (Fig. 1B). This demonstrates that a large portion of the increase in [Ca^2+^]i, which occurs in control C_2_C_12_ myotubes in response to depolarization, is a result of L-type channel activation. Nifedipine also has a strong effect on the rise in [Ca^2+^]i that occurs in response to depolarization in the HERG-expressing myotubes (Fig. 1C). It blocks approximately 60, 73, and 77% of the Ca^2+^ increase that occurs initially (at 20, 30, and 40s, respectively; p<0.05) and then ranges from 70-42% from 50-120s. Although the nifedipine block does not appear as strong over time in the HERG-expressing cells, indeed, when the specific nifedipine-sensitive Ca^2+^ transients are determined and compared (Fig. 1D), there is no statistical difference in the initial nifedipine-sensitive response of the HERG-treated and control myotubes and virtually no difference beyond the initial ∼10 seconds post depolarization. We obtained similar results with the L-type channel blocker nicardipine (data not shown). Thus, although HERG is inducing a greater increase in [Ca^2+^]i in response to depolarization, this effect does not appear to be the result of increased Cav1.1 activity at 48 hours after transduction. We explored potential L-type channel involvement further. *L-Type Current Density.* Myotubes were transduced with either a HERG-encoded or an appropriate control virus and after 48 hours were evaluated for L-type current using whole-cell voltage clamp of myotubes (see Methods). The data demonstrate that HERG had no effect on peak L-type channel Ba^2+^ current density (Fig. 2 A,B) except at only the two most positive voltages (Fig. 2C). *Skeletal Muscle L-Type Channel Gene Expression.* Control and HERG-expressing myotubes were assayed for expression of genes encoding Cav1.1, embryonic Cav1.1 (Cav1.1e), Cav1.2, and Cav1.3 using RT-qPCR. No significant differences were found in expression of any of these genes for up to 60 hours after transduction (Fig. 3 A,B; Cav1.2 and 1.3 data not shown). *Skeletal Muscle L-Type Channel α Subunit Protein Abundance.* Lysates from control and HERG-expressing myotubes were immunoblotted (Fig. 3C) using an antibody specific for the skeletal muscle Cav1.1 channel α subunit and protein signal was detected by chemiluminescence (see Methods). The ODs of the Cav1.1 protein band reveal that HERG had no effect on Cav1.1 protein abundance (Fig. 3D). We find no effect of HERG on L-type current amplitude or Cav1.1 channel α subunit gene expression or protein abundance although HERG does increase myotube [Ca^2+^]i (over controls) in response to depolarization.

**Figure 1.**
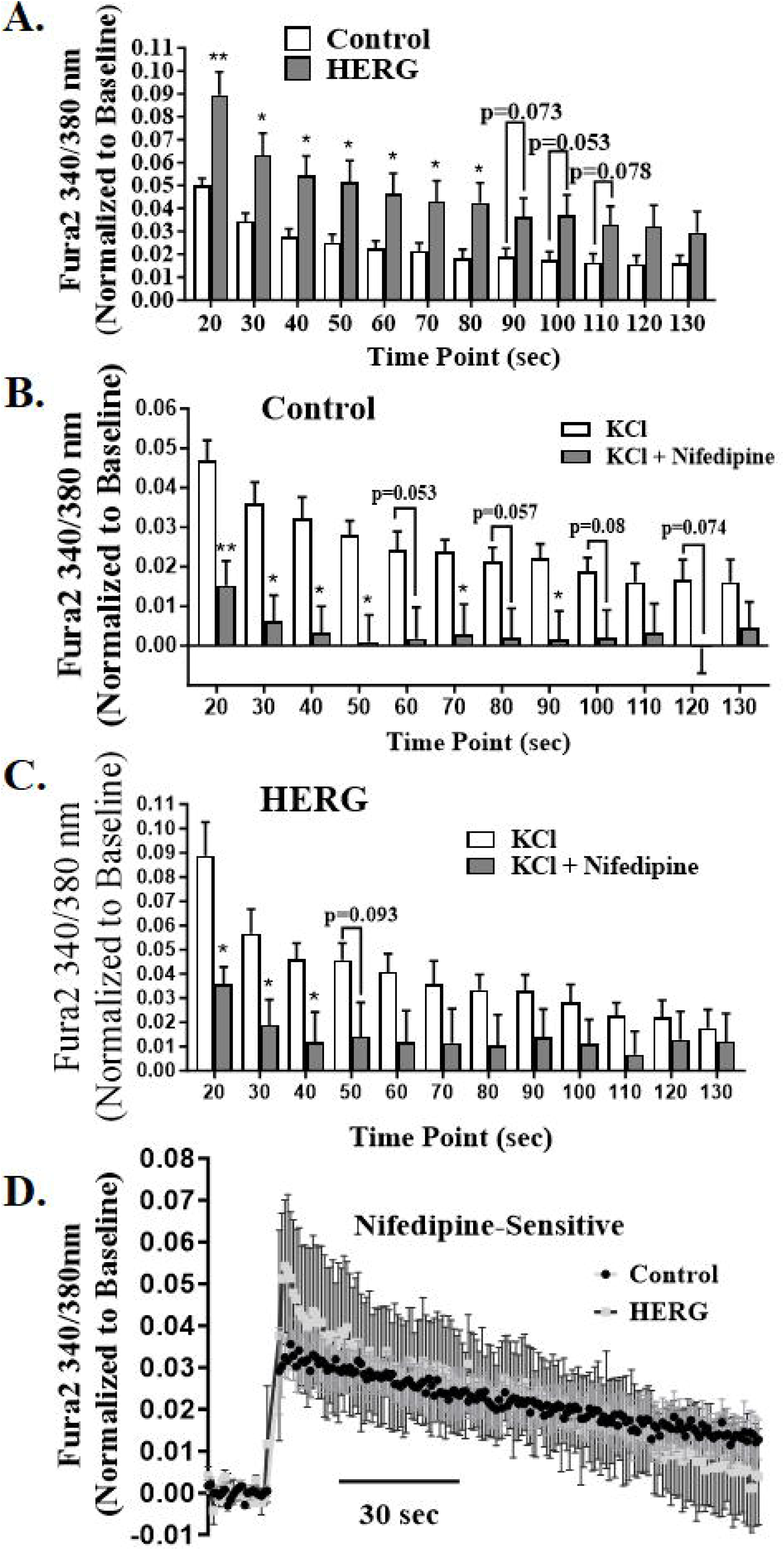
HERG does not enhance intracellular calcium concentration ([Ca^2+^]i) through modulation of the L-type channel in skeletal muscle. A. The increase in intracellular Ca ([Ca^2+^]i) initiated by depolarization (with 100 mM KCl treatment) is significantly greater in myotubes expressing HERG relative to controls. B. Nifedipine (10 μM) has a significant inhibitory effect on the increase in [Ca^2+^]i that occurs in control cells in response to depolarization over 90 seconds after addition of KCl. C. Nifedipine also inhibits a portion of the HERG-induced increase in [Ca^2+^]i for up to 40 seconds. D. Nifedipine-sensitive fluorescence ratios do not differ significantly between the control and HERG-expressing myotubes, demonstrating that the HERG-modulated increase in [Ca^2+^]i does not result from activation of L-type Ca^2+^ channels. [Ca^2+^]i was evaluated by the ratiometric fluorescent Fura-2 dye. The 340/380nm ratios were determined, normalized to baseline, and analyzed by a 2 x 2 ANOVA design for repeated measures. There was no statistically significant interaction between HERG and treatment. The bars (A,B,C) and symbols (D) represent means. The error bars represent standard error of the mean. n=16 (8 control and 8 HERG-expressing myotubes). *p<0.05, **p<0.01.

**Figure 2.**
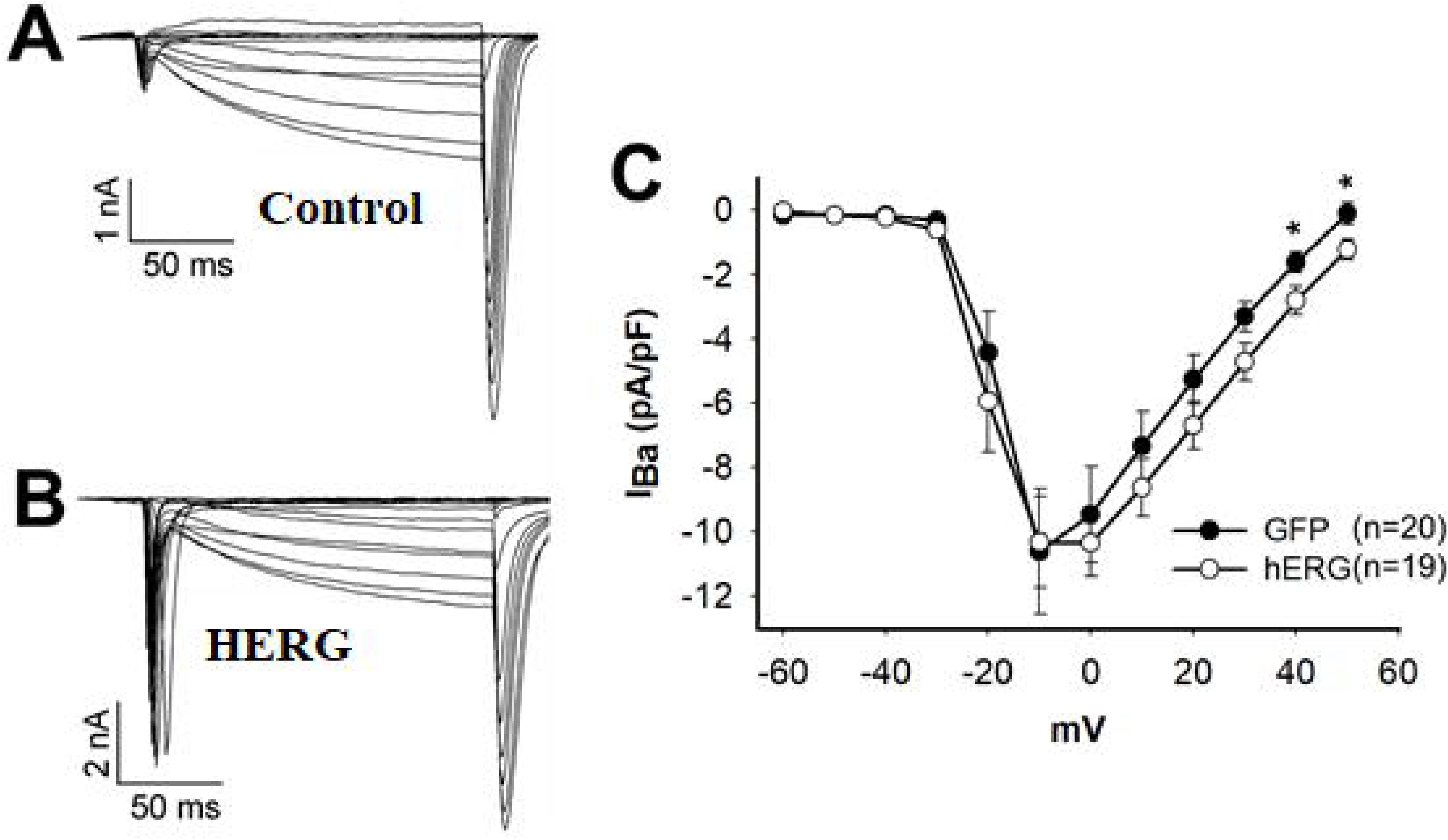
HERG Expression does not change peak Cav channel current density in C_2_C_12_ myotubes. A,B. Example ensembles of current traces elicited by stepping from −80mV to 50 mV in 10 mV increments for 150 msec, from a holding potential of −60 mV in a control transduced myotube A) or a HERG transduced myotube B). Currents in A) and B) were recorded in a bath solution containing 140 mM Na^+^ and 10 mM Ba^2+^ C) Compiled IV curves for peak Ba^2+^(slow) currents measured in control or HERG expressing myotubes. The current density differed only at the two most positive voltages (40 and 50 mV), *p<0.05.

**Figure 3.**
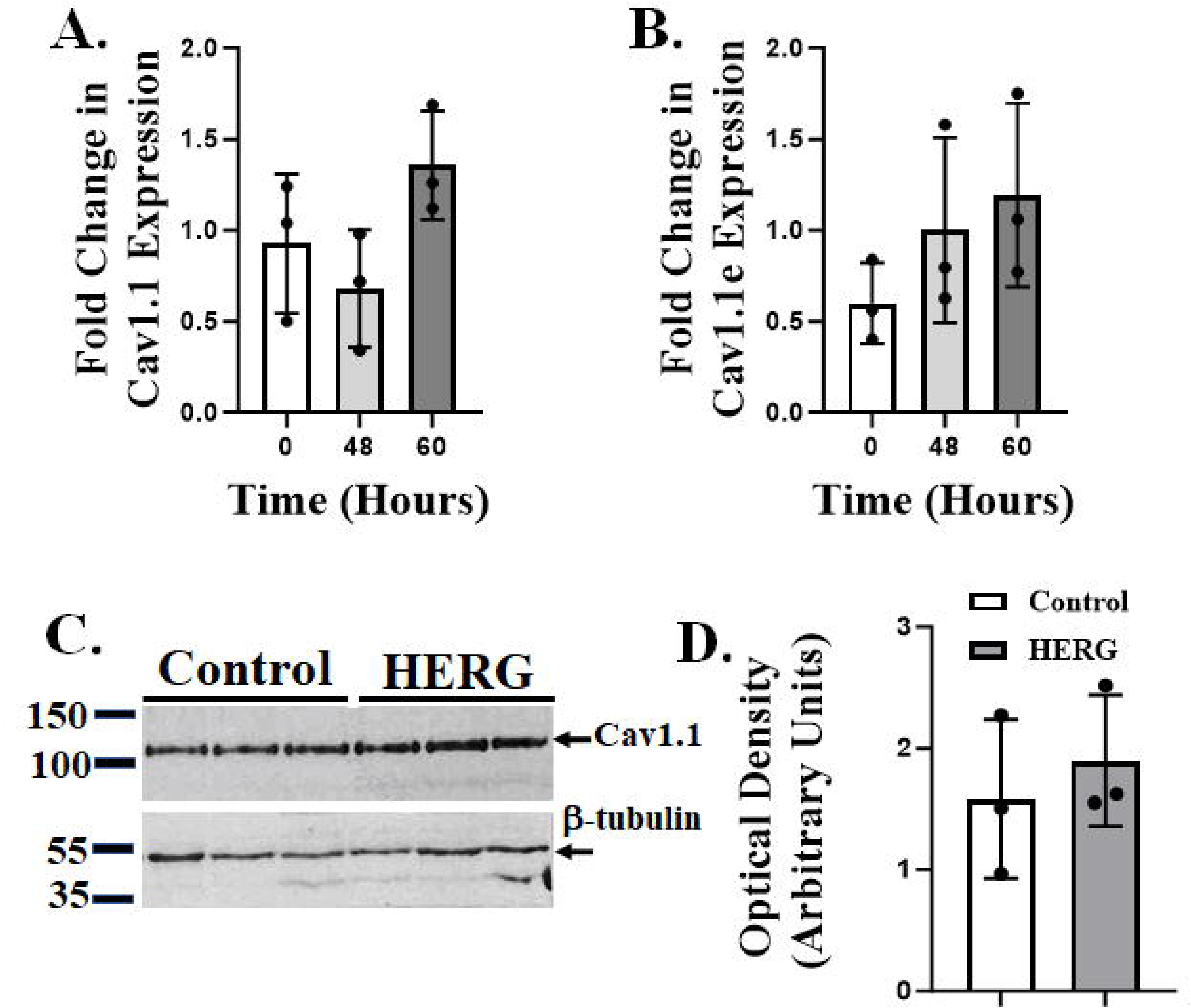
Cav1.1 L-type Ca^2+^ channel gene expression and protein abundance are not affected by HERG expression in C_2_C_12_ myotubes. A,B. rtPCR assays reveal that mRNA levels of adult (A) and embryonic (B) Cav1.1 are not affected by HERG expression up to 60 hours post transduction with HERG encoded adenovirus. *There were no changes noted in levels of Cav1.2 or 1.3 mRNA as well (data not shown). C,D. Immunoblot (C) and optical density measures of protein bands (D) show that Cav1.1 protein abundance is not significantly affected at 48 hours post HERG expression. A,B,D. Bars represent means, error bars represent standard deviations, and filled circles represent a single data point/replicate. A one-way ANOVA was used to analyze the data in panels A and B. Data displayed in panel D were analyzed by a Student’s t-test. A,B. n=9 fold changes) total, derived from 9 control and 9 HERG-expressing samples, divided to produce 3 replicate fold-changes per each of three time points; C,D. n=6, 3 control and 3 HERG-expressing myotube lysate samples. Full blots are available in the Supplementary Data section (SD2).

Indeed, the fact that the HERG-modulated response is activated by depolarization and is nifedipine (10 μM) insensitive implicates ECCE, a Ca^2+^ release pathway that is insensitive to store depletion. *2-APB Block with Depolarization.* To further test if HERG is modulating ECCE, control and HERG-expressing myotubes were treated with vehicle or 2-APB, which will block ECCE (and SOCE) (Olivera & Pizarro 2010), and then depolarized to activate ECCE (and further inhibit SOCE) (Stiber et al., 2008 in Dirksen 2009; Kurebayashi & Ogawa 2001). Time resolved assays of [Ca^2+^]i using Fura-2 revealed that at 15 seconds in control cells, 2-APB inhibits (by 46.9%) the increase in [Ca^2+^]i that occurs in response to depolarization, but the change is not statistically significant (p=0.22; Fig. 4A). However, the initial (up to 15 seconds) increase in [Ca^2+^]i that occurs in response to depolarization in HERG-overexpressing myotubes is inhibited 84.5% (p<0.005; Fig. 4B) and the inhibition continues up to 57 seconds. Indeed, the 2-APB sensitive response during initial treatment (15 s) was significantly greater (75.3%; p<0.05) in the HERG-expressing cells relative to controls (Fig. 4C). Thus, these data suggest that HERG overexpression in C_2_C_12_ myotubes modulates ECCE.

**Figure 4.**
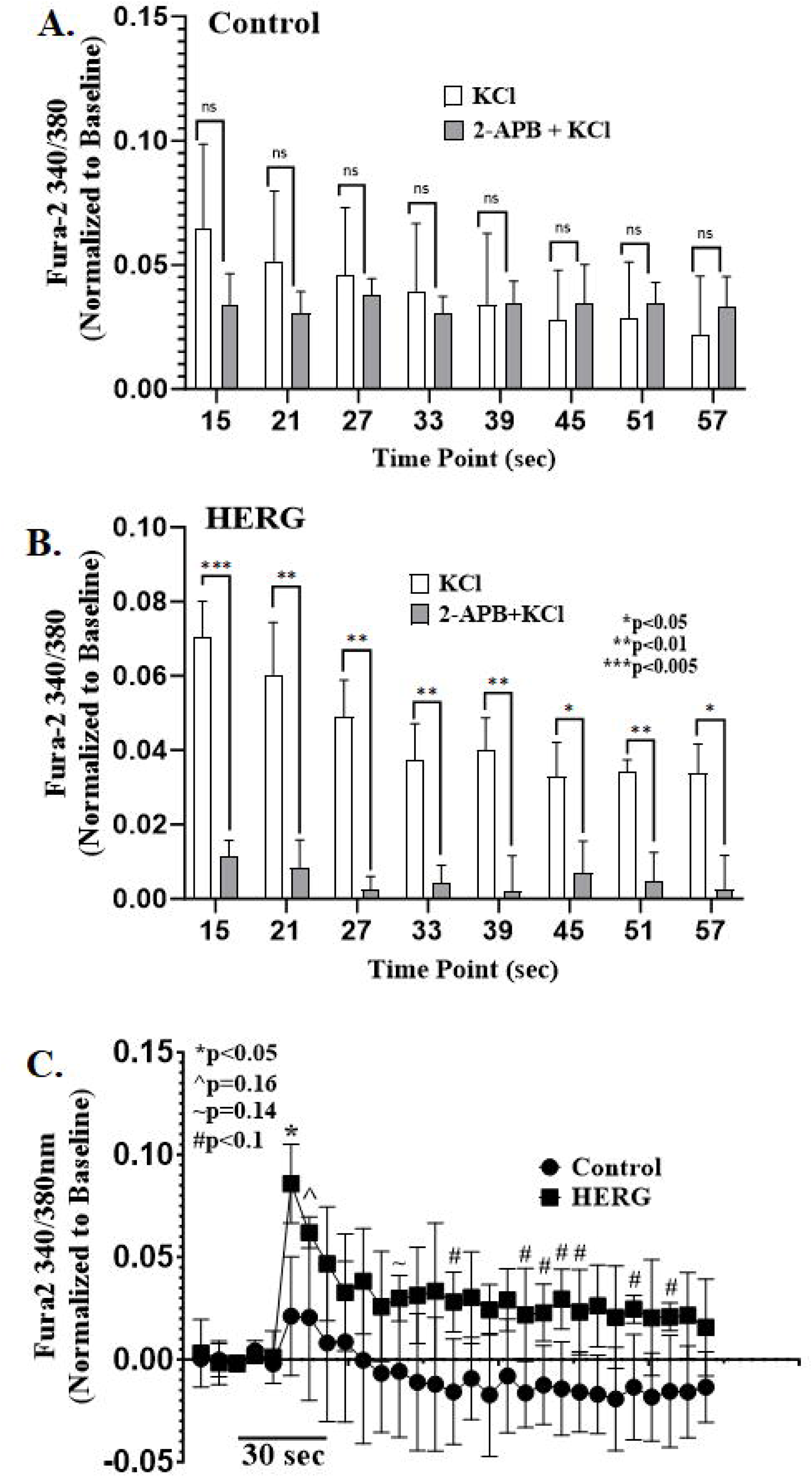
Excitation coupled calcium entry (ECCE) is a source of the greater increase in intracellular calcium concentration ([Ca^2+^]i) that is elicited in response to depolarization by KCl in HERG-expressing cells. A. Initially, in control cells, 2-APB appears to have a mild, but statistically insignificant inhibitory effect on the initial mean increase in [Ca^2+^]i that occurs in response to depolarization. B. The 2-APB has a statistically significant inhibitory effect on the HERG-modulated increase in [Ca^2+^]i that occurs in response to depolarization. C. The initial 2-APB-sensitive currents differ significantly between the depolarized control and HERG-expressing myotubes, suggesting that a source of the initial HERG-modulated increase in [Ca^2+^]i is extracellular. The [Ca^2+^]i was evaluated by the ratiometric fluorescent Fura-2 dye and the 340/380 ratios were determined and normalized to baseline. The data of panels A and B were analyzed by a 2 x 2 ANOVA design for repeated measures; the interaction between HERG and treatment was not statistically significant. For panel C, the mean difference and standard error of the mean difference were used to estimate p values using a comparison of means calculator (see methods). The bars (A,B) and symbols (C) represent means of time frame units and error bars represent standard deviations. n=24 (12 control and 12 HERG-expressing myotubes).

### The HERG channel modulates Ryanodine Receptor 1

#### Thapsigargin Block with Depolarization

Because RyR1 is a component of ECCE activity, we hypothesized that HERG could also be enhancing release of Ca^2+^ from intracellular stores. To test this, control and HERG-expressing myotubes were treated with either vehicle or Tg, which will deplete intracellular stores and activate SOCE, but have no effect on ECCE (Cherednichenko et al., 2004; Lyfenko & Dirksen 2008 in Dirksen 2009). The cells were then depolarized: 1) to release RyR1 from occlusion by Cav1.1 (i.e., DHPR) and enable Ca^2+^ release from the SR to the cytosol through the RyR1 channel (Pitake & Ochs 2015); and 2) to inhibit SOCE (Stiber et al., 2008 in Dirksen 2009; Kurebayashi & Ogawa 2001). Indeed, depolarization activates the ECCE and pre-treatment with Tg will empty SR stores; thus, any inhibition noted in the Tg treated groups relative to vehicle treated groups will be predominantly a consequence of lowered SR stores.

Time resolved [Ca^2+^]i assays using Fura-2 assays revealed thapsigargin did not significantly reduce the rise in [Ca^2+^]i upon KCl depolarization in control cells (Fig. 5A). However, the depolarization-induced rise in [Ca^2+^]i was significantly inhibited by thapsigargin in HERG-expressing myotubes relative to vehicle treated HERG-expressing myotubes for up to 40 seconds after depolarization (20s, 49.4% inhibition, p<0.05; 30s, 72.7%, p<0.05; 40s, 77.1%, p<0.05; Fig. 5B). Indeed, the Tg-sensitive portion of the response to depolarization was greater in HERG-expressing cells, with the difference in the Tg-sensitive response being statistically significant in earlier time points (∼8-14s; Fig. 5C). These data strongly suggest that a source of increased [Ca^2+^]i in the HERG-expressing myotubes is, at least in part, release of Ca^2+^ from SR stores. Thus, we decided to determine if HERG affects RyR1 activity. *Ryanodine Receptor Activation with Caffeine.* Membrane depolarization will uncouple RyR1 from occlusion by the DHPR and allow Ca^2+^ release from the SR to the cytosol through the RyR1 channel (Pitake & Ochs 2015); therefore, we determined if HERG affects RyR1 activity specifically. Thus, RyR1 activity was assayed in control and HERG-expressing myotubes using caffeine to activate the RyR1 and the area under the curve (AUC) of the change in [Ca^2+^]i was determined for each treatment group. (Fig. 6). Caffeine treatment resulted in significant increases in [Ca^2+^]i in both the control and the HERG-overexpressing myotubes and this increase was significantly inhibited by ryanodine (90 M) in both sets. A 2×3 ANOVA was used to compare drug effects on [Ca^2+^]i in response to viral treatment (HERG or control). The data show that, although the interaction between drug and viral treatment was not statistically significant (p=0.1373), there was a statistically significant mean difference between HERG overexpressing and control cells across all drug treatments (p=0.0206). The data demonstrate that HERG overexpression enhances caffeine-induced release of Ca^2+^ from the SR via RyR1 in C_2_C_12_ myotubes.

**Figure 5.**
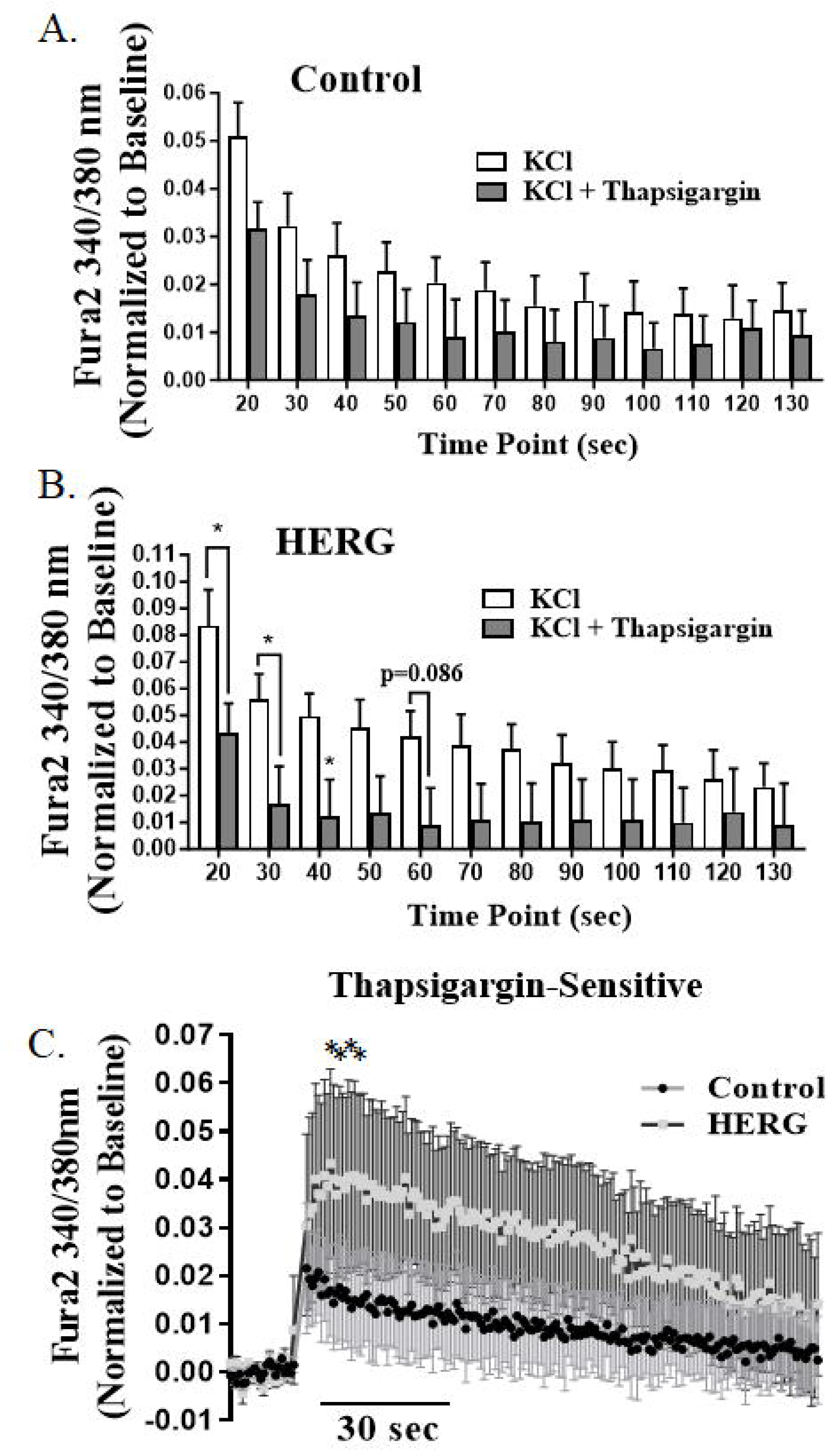
Sarcoplasmic reticulum Ca^2+^ stores is a source of the greater increase in intracellular Ca^2+^ concentration ([Ca^2+^]i) that occurs in response to depolarization by KCl (100 mM) in HERG-expressing cells. A. In control cells, thapsigargin (1 mM) has no statistically significant effect on [Ca^2+^]i in response to depolarization. B. Thapsigargin has a significant inhibitory effect on the HERG-modulated increase in [Ca^2+^]i that occurs in response to depolarization. C. Initial thapsigargin-sensitive fluorescent ratios differ significantly between the depolarized control and HERG-expressing myotubes, suggesting that HERG may also affect release of Ca from intracellular stores. The [Ca^2+^]i was evaluated by the ratiometric fluorescent Fura-2 dye and the 340/380 ratios were determined and normalized to baseline. The fluorescence ratios at noted timepoints (in panels A and B) were analyzed by a 2 x 2 ANOVA design for repeated measures. There was no significant interaction between HERG and treatment. For panel C, the mean difference and standard error of the mean difference were used to estimate p values using a comparison of means calculator (see methods). The bars (A,B) and symbols (C) represent mean fluorescence ratios (i.e., [Ca^2+^]i levels) and error bars represent standard deviation. n=20 (10 control and 10 HERG-expressing myotubes). *p<0.05.

**Figure 6.**
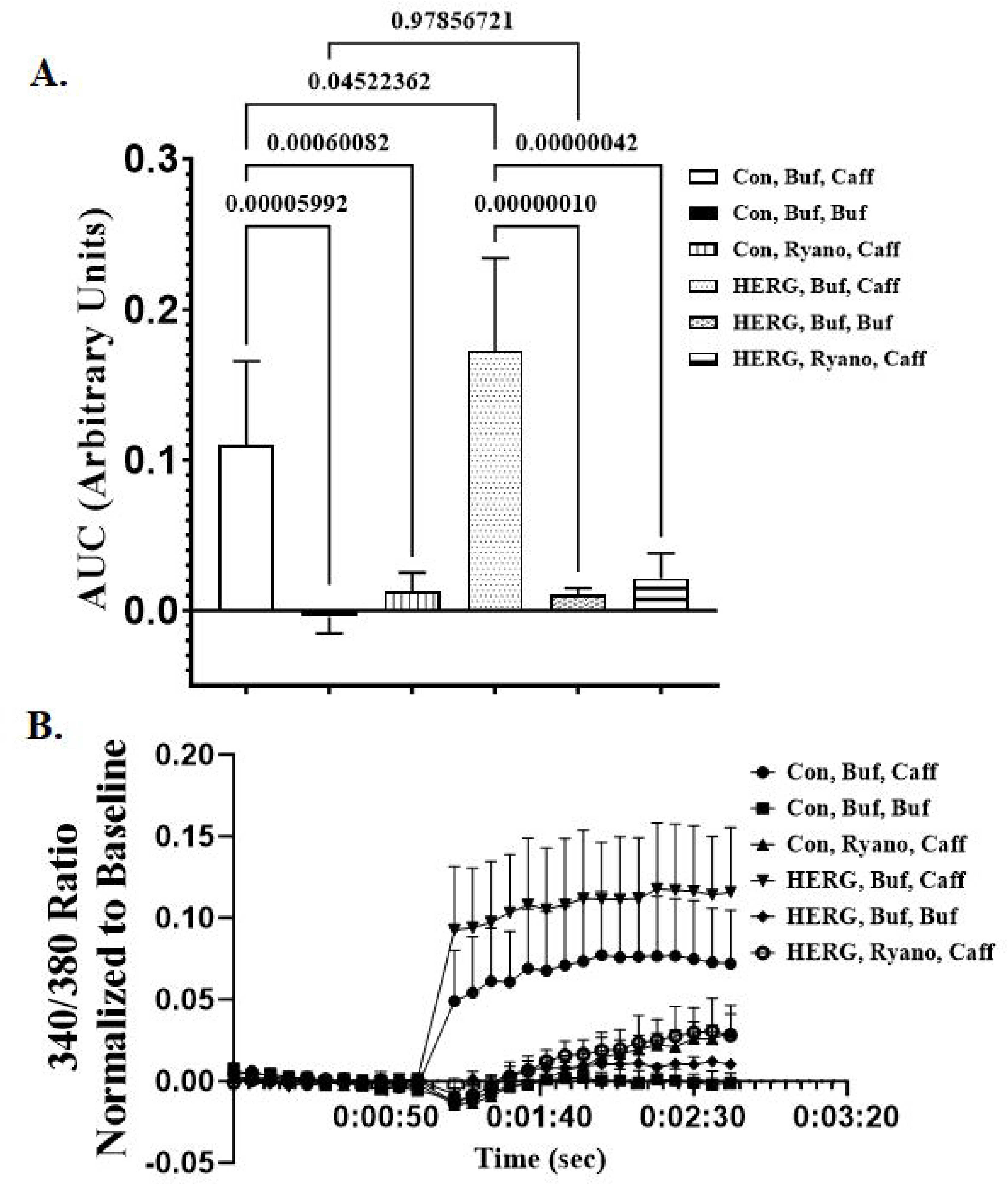
HERG modulates ryanodine receptor-mediated Ca^2+^ release. A. HERG-overexpressing myotubes exhibit a significantly greater increase in intracellular concentration ([Ca^2+^]i) than control cells when treated with caffeine (10 mM); this response is significantly inhibited by ryanodine (90 uM) in both HERG-expressing and control myotubes, but the inhibition is more statistically significant in the HERG-overexpressing cells. n=36 wells (18 wells of HERG-expressing myotubes and 18 wells of controls). B. Representative line graph of a single assay of 24 wells. n=24; the 24 wells consisted of 3 wells per each of six groups. The [Ca^2+^]i was evaluated by ratiometric fluorescent Fura-2 dye and the 340/380 ratios were determined and normalized to baseline. Areas under the curve (AUCs) were analyzed by one-way ANOVA and means were separated by Tukey’s test. The bars and symbols (B) represent means and error bars represent the standard deviation.

### The HERG channel affects [Ca^2+^]i by modulation of Store Operated Calcium Entry

The data strongly support that a source of the HERG-modulated increase in [Ca^2+^]i in skeletal muscle cells is the ECCE; however, this does not rule out the possibility that HERG may also modulate other Ca^2+^ entry pathways such as SOCE. Here, we assay SOCE (see methods) and quench ECCE by not depolarizing the cells. Briefly, we treated control and HERG-expressing cells with Tg to deplete intracellular stores and thus activate SOCE. We then treated cells within each group with either 2-APB to block SOCE (and ECCE) or vehicle. Finally, we added extracellular Ca^2+^ (2.5 mM CaCl_2_) to stimulate SOCE and monitored [Ca^2+^]i with Fura-2. The AUC data show that [Ca^2+^]i in HERG-expressing cells was significantly increased (81%; p<0.04) over controls (Fig. 7) in response to 2.5 mM extracellular CaCl_2_. The 2-APB treatment did not significantly inhibit the rise in [Ca^2+^]i upon addition of 2.5 mM CaCl_2_ in control myotubes. In contrast, 2-APB blocked the [Ca^2+^]i increase that occurred in response to CaCl_2_ in the HERG-expressing cells by a statistically significant 59% (p<0.01). These data demonstrate that overexpression of HERG in C_2_C_12_ myotubes results in a greater SOCE response to thapsigargin, although we do not know if the effect is direct or results from HERG-enhanced depletion of SR stores.

**Figure 7.**
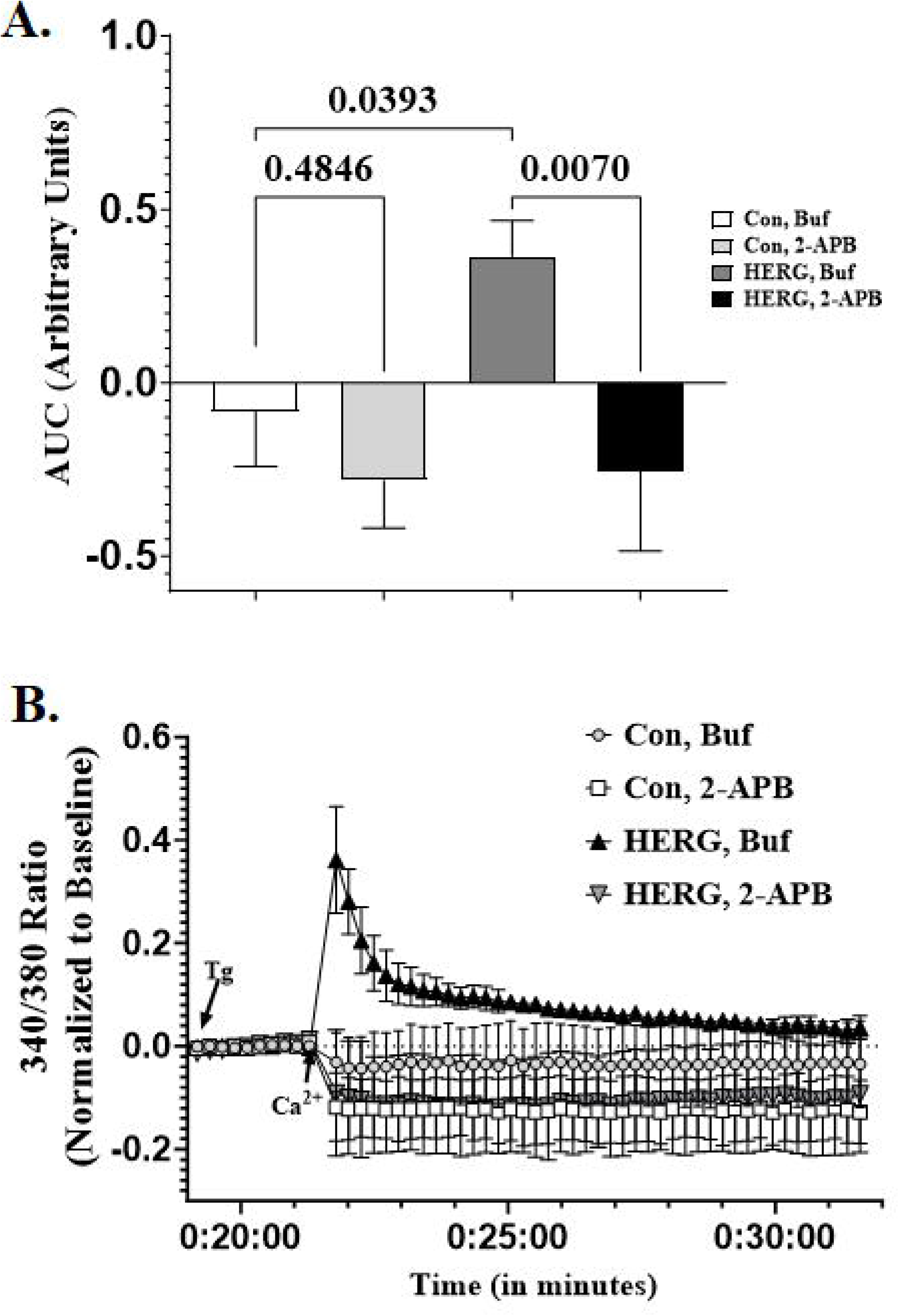
HERG modulates SOCE. A. HERG expressing myotubes exhibit a significantly greater increase in intracellular calcium concentration ([Ca^2+^]i) than control cells when treated with high calcium (2.5 mM) after depletion of SR calcium stores by thapsigargin (Tg, 1 μM) treatment; this response was significantly inhibited by the SOCE inhibitor 2-APB (100 μM). B. Graphic showing the assay over time. The [Ca^2+^]i was evaluated by fluorescent Fura-2 dye: the 340/380 ratios were determined and normalized to baseline. For panel A, the area under the curve (AUC; after addition of calcium) was calculated per treatment group and analyzed by one-way ANOVA. When different, means were separated by Tukey’s test. The bars (A) and symbols represent means and error bars represent the standard deviation. n=12 wells per viral treatment (6 wells of HERG-expressing myotubes and 6 wells of controls; per viral treatment, 3 wells received 2-APB and 3 received vehicle).

### The HERG channel lowers abundance of Calsequestrin1 mRNA and protein

Calsequestrin1 (CaSeq1) functions as both a Ca^2+^ buffer and a Ca^2+^ sensor in the SR and inhibits ryanodine receptor activity at cytosolic Ca^2+^ concentrations around 1 mM in skeletal muscle. It also inhibits the dimerization of the STIM1 protein necessary to activate the Orai1 channel for SOCE activity (Zhang et al., 2016; Jeong et al., 2021). Because the calsequestrin1 protein modulates RyR1 and STIM1, it has the potential to affect numerous pathways by which Ca^2+^ enters the cytosol, including ECC, ECCE, and SOCE. Therefore, we explored the effect of HERG-expression on CaSeq1 levels in C_2_C_12_ myotubes. Total RNA was extracted from both control and HERG-expressing myotubes transduced at 200 MOI (n=8, 4 control plates and 4 HERG plates) and HERG and CaSeq1 mRNA levels, relative to GAPDH, were compared using RT-qPCR. Myotubes transduced with adenovirus encoding HERG exhibited a 2.6-fold increase (p<0.02) in HERG mRNA and a 0.83-fold decrease (p<0.05) in CaSeq1 mRNA at 48 hours after transduction (Fig. 8A). Further, both control and HERG-expressing myotubes were transduced at 200 MOI (n=8, 4 control and 4 HERG) and lysates were immunoblotted using antibodies specific for either CaSeq1 or GAPDH (SD3). We measured the optical densities (OD) of the protein bands and noted a reduction in CaSeq1 protein (23.5%; p=0.081) that approached, but did not reach significance (defined earlier as p<0.05). To increase HERG expression levels, we transduced both control and HERG-expressing myotubes at 400 MOI (n=8, 4 control and 4 HERG) and immunoblotted the lysates (Fig. 8 B-E) with antibodies specific for CaSeq1 and then (after stripping the PVDF membrane) with antibodies specific for GAPDH. We detected a full length CaSeq1 ∼65 kD protein along with ∼50 kD and ∼40 kD CaSeq1 proteins which appear to be CaSeq1 degradation products (Fig. 8B). The mean OD of each CaSeq1 protein band was decreased in the HERG-expressing myotubes relative to control cells: the ∼65 kD protein decreased 89.7% (p=0.012); the ∼50 kD protein decreased 66.3% (p=0.077); and the ∼40 kD protein decreased 80.8% (p=0.088). When the OD values are combined per sample (“Total,” Fig. 8E) and compared by a Student’s t-test (HERG versus control), there is a 77% (p=0.036) decrease in CaSeq1 protein abundance in HERG transduced myotubes. These data show that HERG overexpression produces a decrease in CaSeq1 protein abundance.

**Figure 8.**
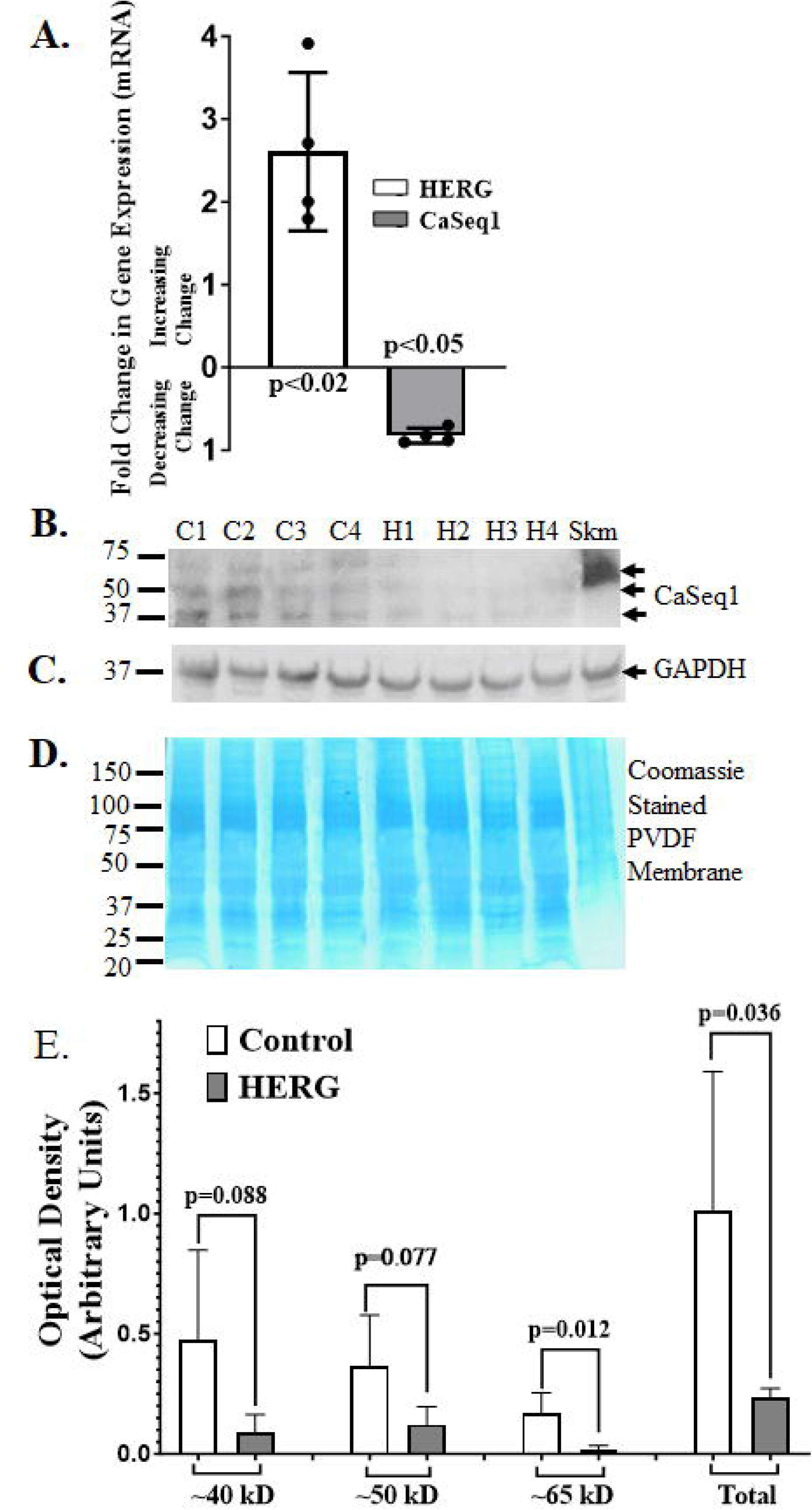
Calsequestrin 1 (CaSeq1) mRNA and protein levels are reduced in C_2_C_12_ myotubes at 48 hours after transduction with 400 MOI HERG encoded virus relative to myotubes transduced with control virus. A. Fold changes in HERG and CaSeq1 mRNA levels in response to transduction with HERG-encoded adenovirus. HERG mRNA increased 2.6-fold (p<0.02) in HERG-transduced myotubes and CaSeq1 mRNA decreased 0.83-fold (p<0.05) in the HERG-expressing cells. B. Control and HERG-expressing myotube lysates immunoblotted with antibody specific for CaSeq1 protein. C. Control and HERG-expressing myotube lysates immunoblotted with antibody specific for the “house-keeping” protein GAPDH. D. PVDF membrane stained with Coomassie blue to confirm equal sample loading in lanes. E. Normalized optical densities (OD) of individual CaSeq1 proteins and of CaSeq1 proteins combined (“Total”) from immunoblot. Mean ODs of control and HERG-expressing cells (within each protein) were analyzed by Student’s T-test. Bars represent average OD and error bars denote the standard deviation. n=8, 4 control groups and 4 HERG-expressing groups. Skm = mouse skeletal muscle. Full blots are available in the Supplementary Data section (SD3).

**Figure 9.**
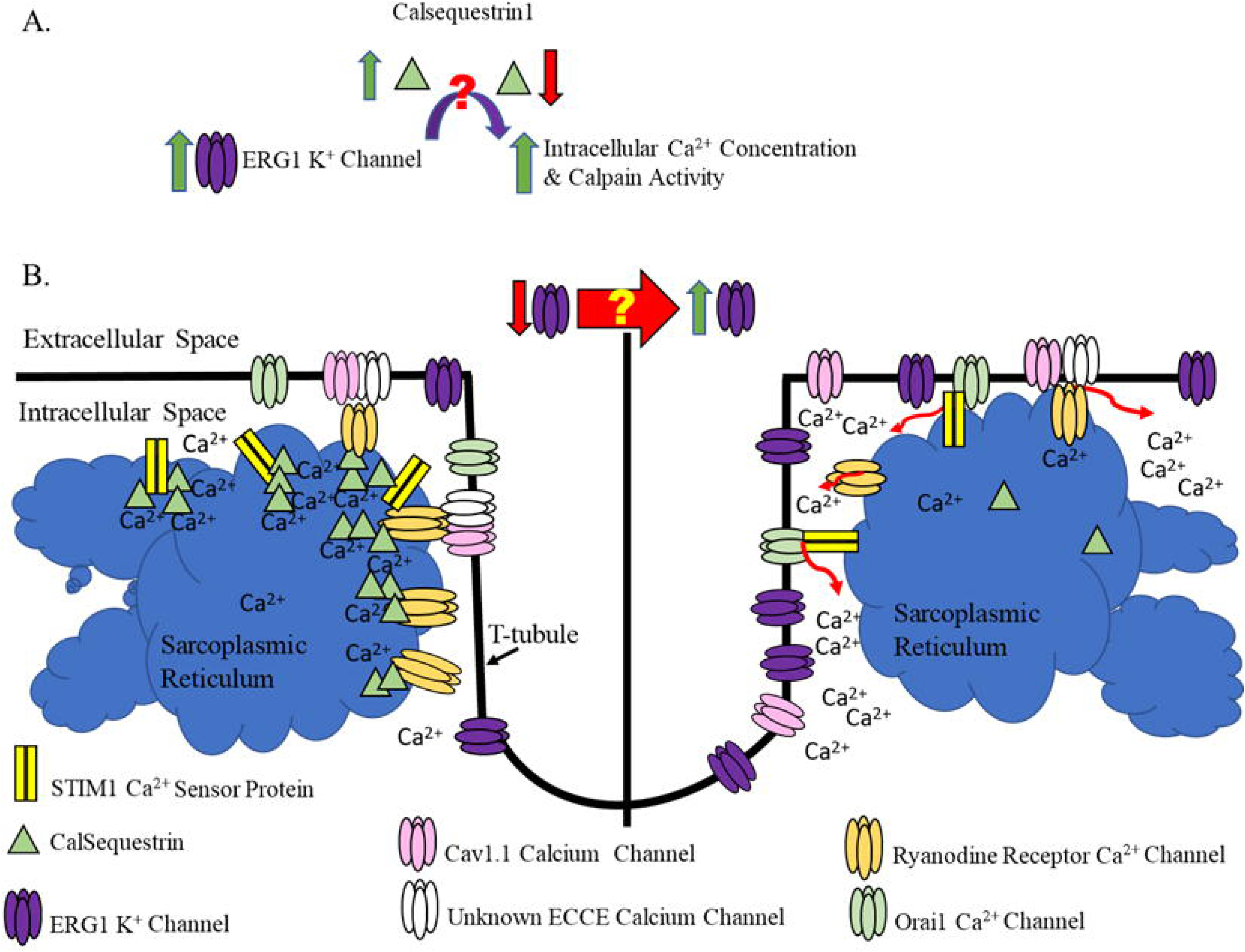
HERG increases intracellular Ca^2+^ concentration by decreasing abundance of the Ca^2+^ binding/buffering protein Calsequestrin1. Two important questions are: A) How does ERG1A decrease Calsequestrin1?; and B) What induces increased levels of the ERG1A channel itself?

## DISCUSSION

Although ERG1A has been detected in the sarcolemma and t-tubules of heart muscle (Rasmussen et al., 2004; Jones et al., 2004), the presence of ERG1A in skeletal muscle was not reported until we detected it in the sarcolemma of atrophying skeletal muscle (Wang et al., 2006; Zampieri et al., 2021). We reported that overexpression of ERG1A in mouse *Gastrocnemius* muscle enhances proteolysis through up-regulation of UPP activity by increasing the protein (and mRNA) abundance(s) of the muscle specific UPP E3 ligase MuRF1 (Wang et al., 2006; Pond et al., 2013; Hockerman et al., 2014). Further, we showed that HERG overexpression in C_2_C_12_ myotubes also produces an increase in MuRF1 protein abundance and, interestingly, in both basal [Ca^2+^]i and calpain activity (Whitmore et al., 2020). Because elevated [Ca^2+^]i is known to play a role in atrophy (Sartori et al., 2021), we sought to determine by what mechanism(s) HERG regulates [Ca^2+^]i.

ERG1A is a sarcolemmal membrane-bound voltage-gated K^+^ channel, thus it is feasible that it might in some way affect the L-type voltage-gated Ca^2+^ channel(s) also located in the sarcolemma. We reasoned that the ERG1A channel would not likely hyperpolarize the skeletal myocyte membrane appreciably because it’s role in cardiac tissue is to facilitate repolarization of the action potential and prevent early after depolarizations (Jones et al., 2014). We hypothesized, however, that the channel’s presence in the skeletal muscle sarcolemma might in some way modulate the current amplitude or abundance of the Cav1.1 Ca^2+^ channels (DHPRs) also located there. Indeed, the depolarization-induced calcium flux in the HERG overexpressing cells was greater than in control cells; however, the difference between the KCl-stimulated Ca^2+^ increase in control and the hERG overexpressing cells is not significantly inhibited by nifedipine (10 μM) (or nicardipine 10 μM, data not shown) and thus cannot be accounted for by L-type channel mediated Ca^2+^ influx. Thus, we postulated that it could result from activation of the ECCE. Indeed, the ECCE is an extracellular Ca^2+^ entry pathway that is activated by prolonged and repetitive depolarization (e.g., 100 mM KCl), but is not sensitive to block by 10 μM nifedipine (Bannister et al., 2009). Block of ECCE activity in primary mouse muscle cultures requires a higher concentration of nifedipine (50 μM, Bannister et al., 2009). Thus, to test for possible HERG modulation of ECCE, we applied 2-APB (known to block ECCE activity; Cherednichenko et al., 2004), and discovered that the HERG-modulated, depolarization-induced increase in [Ca^2+^]i is inhibited by 2-APB. SOCE, which allows calcium entry from the extracellular milieu, is also blocked here by both depolarization and 2-APB (Cherednichenko et al., 2004). Further, the assay conditions did not deplete the SR, which will also activate SOCE (Kurebayashi & Ogawa 2001; Stiber et al., 2008). Therefore, here we have evidence that HERG modulates [Ca^2+^]i (at least in part) through modulation of ECCE.

The ECCE activity unit is believed to be composed of Cav1.1, RyR1, and possibly an (as yet) unrevealed calcium conducting and/or modulating moiety (Bannister & Beam 2013; Cho et al., 2017; Friedrich et al., 2010; Dirksen 2009). Because of the involvement of RyR1 in ECCE activity, we hypothesized that HERG might also affect release of calcium from intracellular stores through RyR1. To test this, we investigated the effect of SR depletion (induced by Tg inhibition of SR Ca^2+^-ATPase) on the HERG-modulated elevation of [Ca^2+^]i. Indeed, we discovered that (at least a portion of) the HERG-modulated, depolarization-induced increase in [Ca^2+^]i is sensitive to store depletion by Tg, suggesting that HERG may also activate release of Ca^2+^ from intracellular stores (Cherednichenko et al., 2004; Hurne et al., 2005). To explore this further, we investigated the effect of HERG on RyR1 activity more directly. RyR1 is a caffeine activated, voltage-gated protein (Nelson et al., 2013) which will allow calcium release from the SR (Pitake & Ochs 2015). Here we demonstrate that HERG overexpression enhances the increase in [Ca^2+^]i that occurs with caffeine treatment in myotubes; this effect is blocked by inhibitory concentrations of ryanodine, suggesting that HERG modulates RyR1 activity. Our data also suggest that HERG modulates SOCE; however, it is not clear whether this HERG-induced increase in SOCE activity is a consequence of stores depletion subsequent to RyR1 activation OR of direct modulation of SOCE pathway components. This remains to be explored.

Our data confirm that HERG overexpression affects [Ca^2+^]i in cultured myotubes. Further, it strongly suggests that it does so (at least in part) by enhancing RyR1 activity, a known component of the ECCE. Indeed, we also show that ECCE activity is enhanced by HERG overexpression. However, our data also show that HERG does not affect myotube L-type current amplitude over time nor does it modulate Cav1.1 protein abundance nor mRNA levels. These findings could suggest that an L-type channel (specifically Cav1.1) is not part of ECCE activity OR that, if it is, then pore permeation of this channel by calcium is not necessary for ECCE activity (as is suggested by earlier work) (Cherednichenko et al., 2004). Nonetheless, it has been shown that the characteristics of ECCE calcium flux are those of L-type currents and that presence of Cav1.1 L-type channels are necessary for ECCE (Bannister et al., 2009; Cherednichenko et al., 2004). Interestingly, however, one report investigating calcium flux in mouse skeletal muscle fibers describes a voltage-activated Mn^2+^ pathway which is “parallel to,” but distinct from, the L-type voltage-activated calcium channel and has properties which suggest it could contribute to ECCE activity (Berbey & Allard 2009). Perhaps this “parallel” entity may be a more likely candidate for current conduction in the ECCE.

Because our data suggest that ERG1A modulates both RyR1 and ECCE activities, as well as SOCE activity, we conjectured that ERG1A could be affecting an entity that modulates all three Ca^2+^ entry pathways. We hypothesized that CaSeq1, a Ca^2+^ sensing and buffering protein detected prominently within the SR milieu, would be a likely candidate. Indeed, our immunoblot data demonstrate that CaSeq1 protein abundance is significantly lower in HERG-expressing cells than in controls. Interestingly, one known mechanism by which CaSeq1 modulates [Ca^2+^]i is by interacting with RyR1. At ≥5 mM Ca^2+^ concentration in the SR ([Ca^2+^]_SR_), CaSeq1 is polymerized and binds/stores high Ca^2+^ levels near the junctional SR membrane to allow quick release of Ca^2+^ during excitation contraction coupling; this action also buffers Ca^2+^ and regulates SR osmolarity (Perni et al., 2013; reviewed in Woo et al., 2020 and in Wang & Michalak 2020). At ∼1 mM [Ca^2+^]_SR_, although still polymerized and binding Ca^2+^ ions, CaSeq1 binds and inhibits RyR1 so that Ca^2+^ cannot pass through this channel into the cytosol (Beard et al. 2005; Herzog et al., 2000). At [Ca^2+^]_SR_ ≤100 μM, CaSeq1 depolymerizes, thus lowering its Ca^2+^ binding capacity and releasing free Ca^2+^ within the SR as it also dissociates from RyR1. The interaction between RyR1 and CaSeq1 requires other SR junctional proteins such as junctin and triadin (Wang & Michalak 2020). Indeed, removing CaSeq1 from the RyR1-junctin-triadin complex creates an increased probability and duration of RyR1 channel opening; adding the CaSeq1 back to this RyR1 complex yields decreased channel opening duration [(Beard et al., 2002; Beard et al., 2005]). Thus, CaSeq1 essentially blocks RyR1 activity at resting Ca^2+^ concentrations.

Further, RyR1 is a component of ECCE and, although it is not believed to be the Ca^2+^-passing entity in this extracellular Ca^2+^ entry pathway (reviewed in Dirksen 2009 and Cho et al., 2017), ECCE activity is modulated by ryanodine treatment: accentuated by low dose and blocked by high doses as is the RyR1 protein (Gach et al., 2008; Cherednichenko et al., 2004, 2008). Thus, it is possible that our observed effect of ERG1A on ECCE is through the lowered CaSeq1 abundance and consequent decreased RyR1 activity. Although less likely, it could also be through CaSeq1 modulation of Cav1.1. Indeed, a complex of CaSeq1 and the transmembrane protein JP45 modulate Cav1.1 (Mosca et al., 2013) and ablation of both JP45 and CaSeq1 proteins enhances ECCE activity in mouse skeletal muscle (Mosca et al., 2016). However, as we demonstrate here, ERG1A modulation of Cav1.1 current conduction is not likely to occur. However, ERG1A could also be affecting [Ca^2+^]i through direct CaSeq1 modulation of SOCE. CaSeq1 regulates SOCE by inhibiting STIM1 aggregation, thus decreasing the entry of Ca^2+^ through this extracellular pathway (Wang et al., 2015; Zhang et al., 2016). Thus, lowered CaSeq1 abundance could result in less inhibition of STIM1 aggregation and increased [Ca^2+^]i.

Indeed, down-regulation of CaSeq1 protein abundance could increase [Ca^2+^]i through interaction of multiple mechanisms and play a role in the skeletal muscle atrophy induced by ERG1A. Work with CaSeq1 knockout mice supports this idea. CaSeq1 knockout mice were viable, but skeletal muscle from these mice exhibited enhanced sensitivity to caffeine, an increased magnitude of Ca^2+^ release in response to depolarization, and an increase in resting [Ca^2+^]i, likely because CaSeq1 modulates RyR1 activity (Dainese et al., 2009). Thus, these data support the idea that a decrease in CaSeq1 would result in an increase in both resting [Ca^2+^]i and Ca^2+^ release in response to depolarization. Further, CaSeq1 ablation results in formation of SR-stacks which contribute to formation of Ca^2+^ entry units (CEUs), which are intracellular junctions that modulate SOCE (Dainese et al., 2009). These SR-t-tubule junctions within the I-band of CaSeq1 null mice muscle represent pre-assembled CEUs that provide a mechanism for constitutively active SOCE, hypothesized to be a calcium source compensating for the lower Ca^2+^ concentration within the SR (Boncompagni et al. 2012; 2018; Michelli 2019, 2020).

Constitutively active SOCE, whether induced by direct HERG action on pathway components or through HERG-induced lowered SR calcium stores (resulting from enhanced RyR1 activity) could explain the increased basal intracellular Ca^2+^ detected in C_2_C_12_ myotubes overexpressing HERG (Whitmore et al., 2020). Interestingly, CaSeq1-ablated mice also presented with a reduction in body weight compared to wild type (WT) (Paolini et al., 2015; Paolini et al., 2007) and with skeletal muscle atrophy, which manifested as a significant decrease in both EDL muscle fiber cross sectional area and grip strength (Paolini et al., 2015). The atrophy likely also results from the increase in expression of certain atrogenes (i.e., CathepsinL, Psmd1, Bnip3, and Atrogin) also reported by this group. Additionally, it could be (at least in part) a result of an increase in Ca^2+^ dependent calpain activity induced by the increase in [Ca^2+^]i. These results are consistent with the finding that HERG overexpression in *Gastrocnemius* muscle and in C_2_C_12_ myotubes induces skeletal muscle atrophy (Whitmore et al., 2020; Wang et al., 2006).

An important question left unanswered by this study is: How does ERG1A overexpression result in lowered CaSeq1 protein levels (Fig.8A)? Interestingly, our RTPCR data reveal that ERG1A overexpression decreases CaSeq1 gene expression. Indeed, lowered levels of CaSeq1 mRNA are likely to result in a lower abundance of the encoded protein (as observed here); however, how the large membrane bound HERG protein affects transcription of CaSeq1 will require study. We hypothesize this modulation likely occurs by HERG interaction with small signaling (perchance DNA-binding) proteins that interact with regions of the HERG N-terminus, specifically the PAS-PAC sequence. HERG may also affect CaSeq1 mRNA levels through post-transcriptional modulation. Additionally, we have shown that ERG1A up-regulates UPP activity by increasing the abundance of the MuRF1 E3 ligase in mouse skeletal muscle and in C_2_C_12_ myotubes (Whitmore et al., 2020; Wang et al., 2006). Therefore, we hypothesize that ERG1A could also enhance the degradation of the CaSeq1 protein by increasing UPP activity; indeed, there are ubiquitinylation sites on CaSeq1 (https://www.phosphosite.org/homeAction). These areas remain to be explored.

Perhaps the most fundamental question regarding the role of ERG1A in skeletal muscle atrophy is: “What induces increased levels of the ERG1A channel in skeletal muscle (Fig. 8B)?” ERG1A is up-regulated in skeletal muscle undergoing atrophy as a consequence of unloading, cancer cachexia and denervation (Zampieri et al., 2021; Wang et al., 2006; Hockerman et al., 2013; Pond et al., 2014). It is likely, therefore, that some factor common to these conditions induces expression of ERG1A. Perhaps simply a decrease in stimulation of the muscle contributes in some way. This is another area that remains to be explored.

An important limitation of this study is that all of the work presented here has been performed in cultured myotubes rather than in animals or primary isolated skeletal muscle fibers or tissue explants, which would provide a more holistic view. Indeed, muscle contraction, force generation, metabolic function, etc. are highly affected by/dependent upon specific tissue structural characteristics and the interaction of various cells. The cultured myotubes do not have complete contractile apparati and lack access to extracellular matrix materials and ultrastructure which help localize growth factors and glycoproteins necessary for growth and tissue metabolism (reviewed in Smith & Meyer 2019; Gillies and Lieber 2011). However, the isolated, cultured cells allow for study of skeletal muscle cell physiology without interference from other tissue type effects and, thus, yield a more accurate model for study of [Ca^2+^]i in skeletal muscle cells specifically. Other limitations with this work include the use of an adenoviral vector to overexpress the HERG gene. Although we do control for general viral perturbations by comparing our HERG overexpressing myotubes to control myotubes transduced with the same vector not encoding HERG, we still are using an “overexpression” model. That is, the abundance of HERG protein produced in response to transduction, as well as the cellular response to the HERG protein, may or may not be realistic in terms of how muscle cells respond *in vivo* to certain atrophic stimuli and the subsequent upregulation of HERG. However, we have shown that transduction of C_2_C_12_ myotubes with HERG results in decreased myotube area and increased MuRF1 E3 ligase protein abundance as occurs when ERG1A is electro-transferred into mouse skeletal muscle (Whitmore et al. 2020; Pond et al., 2013; Wang et al., 2006). As well, we have shown that overexpression of HERG in C_2_C_12_ myotubes yields an increase in both [Ca^2+^]i and subsequent calpain activity (Whitmore et al., 2020). Indeed, decreased myofiber size and increased UPP E3 ligase protein abundances as well as increased [Ca^2+^]i and calpain activity are known characteristics of atrophic skeletal muscle (Sartori et al., 2021). Thus, despite these potential limitations, this study has relevance to muscle health and muscle pathologies because perturbances (e.g., increases) in [Ca^2+^]i can have detrimental and even severe effects on muscle tissue (Sartori et al., 2021; Lopez et al., 2018; Goswami et al., 2015).

In summary, this study provides a potential mechanism to explain how upregulation of ERG1A might contribute to increased [Ca^2+^]i and, thus, atrophy in skeletal muscle. Here, we describe downregulation of CaSeq1 protein (and mRNA) abundance in response to HERG overexpression. We also report amplification of Ca^2+^ release from the SR via RyR1 in addition to increased ECCE and SOCE activities in myotubes overexpressing HERG. It is likely that HERG enhancement of RyR1 activity, through decreased CaSeq1 abundance, is increasing [Ca^2+^]i. We suggest that the HERG-modulated effect on the ECCE occurs through modulation of RyR1. We further postulate that HERG overexpression may constitutively enhance SOCE activity perhaps by lowering the concentration/availability of free calcium in the SR and/or by the decreased inhibition of STIM1 resulting from lowered CaSeq1. A greater understanding of the mechanism(s) by which this K^+^ channel affects [Ca^2+^]i could identify suitable targets for development of therapies for atrophy and numerous other skeletal muscle pathologies. Indeed, because ERG1A is a component of the I_Kr_ channel responsible for repolarization of the cardiac action potential, it is not a tractable pharmacological target for treatment of atrophy. Finally, because ERG1A channels are also found in other tissues (e.g., brain, islet cells, tumor cells, heart, etc.) (Hardy et al., 2009; Pond et al., 2000; Perez-Neut et al., 2015), research of this mechanism may ultimately invite a broader interest, addressing ERG1A-modulated Ca^2+^ dysregulation in other tissues (Brawek and Garaschuk, 2014; Cheng et al., 2015; Sama and Norris 2013; Wang et al., 2014).

## ACKNOWLEDGEMENTS

The research reported in this publication was supported in large part by the Department of Defense office of the Congressionally Directed Medical Research Programs through a Peer Reviewed Medical Research Program (PRMRP) Discovery Award to ALP and GHH. The Southern Illinois University Carbondale (SIUC) Graduate School provided some financial support through funding to graduate students CW and SG. SIUC also provided some support through an Undergraduate Research-Enriched Academic Challenge (REACH) Award to OK. The Southern Illinois School of Medicine provided some financial support through their Medicine Mentor Professional Enrichment Experience Program to NM.

