## Supplementary figures and images for "ERG1A K^+^ Channel Increases Intracellular Calcium Concentration through Modulation of Calsequestrin1 in C_2_C_12_ Myotubes"

### Figure SD1.tif

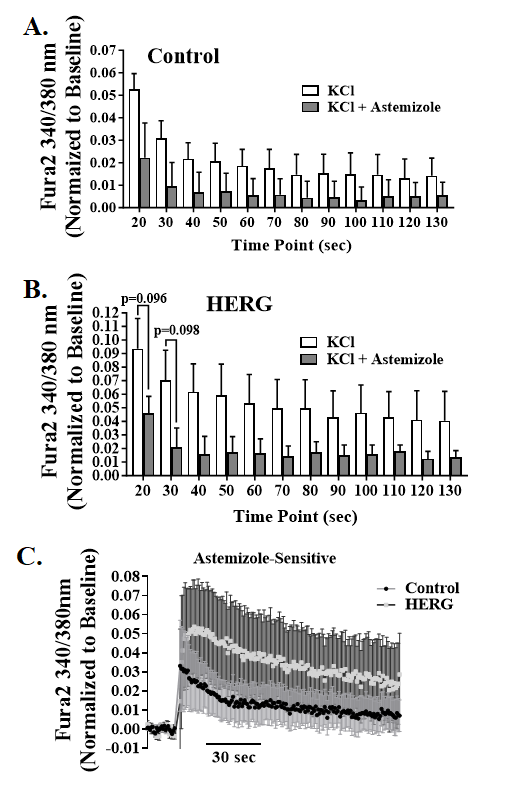

### Figure SD2.tif

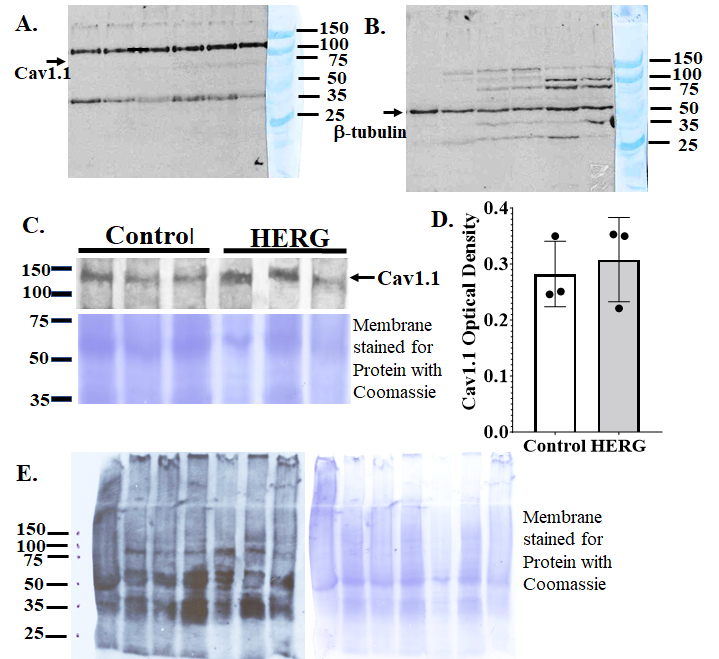

### Figure SD3.tif

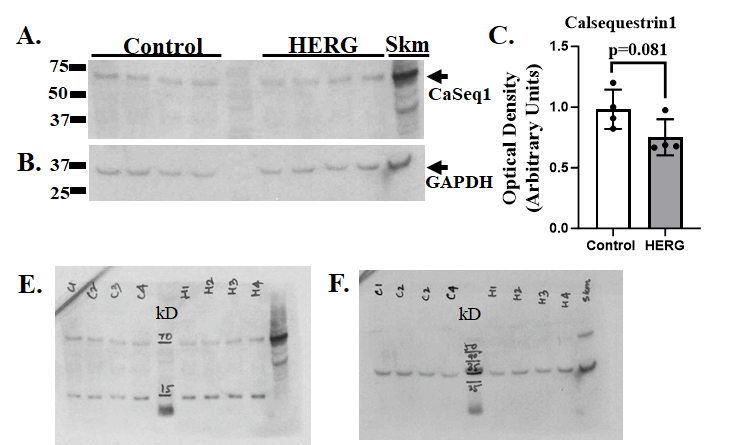
