## Supplemental Figure Legends for "ERG1A K^+^ Channel Increases Intracellular Calcium Concentration through Modulation of Calsequestrin1 in C_2_C_12_ Myotubes"

*Figure SD1. HERG Block by Astemizole.* A. Although the mean $[Ca^{2+}]_i$ is consistently lower in the depolarized control myotubes treated with astemizole ( 1 nM) relative to those treated with vehicle alone over time, the effect of astemizole is not statistically significant. B. The mean $[Ca^{2+}]_i$ is consistently lower in the depolarized HERG-expressing myotubes treated with astemizole relative to those treated solely with vehicle, only *approaching* statistical significance (defined in methods as $p<0.05$) at initial points (20s, 30s). C. The mean astemizole-sensitive response over time is consistently lower in the control cells than in the HERG-expressing cells; however, again the effect is not statistically significant. The $[Ca^{2+}]_i$ was evaluated by the ratiometric fluorescent Fura-2 dye and the 340/380 ratios were determined and normalized to baseline (see Methods). The 340/380 nm ratios determined for indicated timepoints were analyzed by a 2 x 2 ANOVA design for repeated measures. The analysis indicated there was no significant interaction between HERG and treatment. The bars (A, B) and symbols (C) represent means and error bars represent standard error of the mean. n=20 (10 control and 10 HERG-expressing myotubes).

*Figure SD2. HERG over-expression in C2C12 myotubes does not affect Cav1.1 protein abundance.* A,B. These are the original full length immunoblots from which the Cav1.1 (A) and β-tubulin (B, housekeeping protein) blot segments are taken for Figure 3C (40 μg protein per lane). The Cav1.1 blot has a protein at about 35 kD, which may be a low mass degradation product. The β-tubulin blot shows multiple inconsistent, generally faint protein bands in most lanes. These may represent aggregated tubulin where above 50 kD or degradation products where lower. C,D. A previously immunoblotted (C) separate set of samples (60 μg protein per lane) corroborates that HERG over-expression does not affect Cav1.1 protein abundance. Optical density measures (D) of this blot confirm that Cav1.1 protein abundance is not significantly affected at 48 hours post HERG expression. Bars represent means, error bars represent standard deviations. Data were analyzed using a Student's t-test and no real differences were found. n=6 replicates (3 control sets and 3 sets of HERG-expressing myotubes). E. Original blot (left) for Cav1.1 protein (60 μg protein per lane) and the same PVDF membrane stained with Coomassie (right) after blotting. To improve blot clarity, less protein (40 μg) was loaded into each lane for the Cav1.1 immunoblot presented in Figure 3.

*Figure SD3. HERG treatment induces a decrease in Calsequestrin1 (CaSeq1) protein.* A. Immunoblot of control and HERG-expressing myotubes immunoblotted with antibody specific for calsequestrin 1 (CaSeq1). These samples are different than those immunoblotted for CaSeq1 in Figure 7; these myotubes were transduced with 200 MOI adenovirus rather than 400 MOI. Here, CaSeq1 protein is reduced 23.5% (p=0.081) in $C_2C_{12}$ myotubes 48 hours after transduction with HERG encoded virus relative to myotubes treated with control virus. B. Control and HERG-expressing myotubes immunoblotted with antibody specific for GAPDH show that lanes were loaded (40 μg protein) evenly. C. Optical densities (ODs) of CaSeq1 and GAPDH proteins were determined for both control and HERG-expressing cells and normalized to background. A ratio of the normalized ODs (CaSeq1 to GAPDH) was calculated for each sample and these ratios were analyzed by a Student's T-test. Bars represent average OD and error bars denote the standard deviation. n=8, 4 control groups and 4 HERG-expressing groups. Skm = mouse skeletal muscle. Methods. PVDF membranes were immunoblotted with CaSeq1-specific primary antibody (26665-1-AP ProteinTech; Rosemont, IL) in 0.2% non-fat dry milk in Tris-buffered saline with 0.1% tween-20, 5% NGS and 0.1% sodium-azide (pH 7.4, TTBS) followed by washing in TTBS and then incubation in secondary antibody (goat anti-rabbit IgG-Alkaline Phosphatase; 65-6122 Sigma) diluted 1:1000 in 0.2% non-fat dry milk in TTBS. The PVDF membrane was then stripped with stripping buffer (62.5mM Tris Cl [pH=6.8], 2% SDS, 0.695% BME in water) and re-probed with the GAPDH antibody Clone 6C5 (MilliporeSigma; St. Louis, MO) diluted 1:8000 in 0.2% non-fat dry milk in TTBS with 5% NGS and 0.1% sodium azide buffer. The chemiluminescent signal was developed using an ImmunStar AP Western Chemiluminescent Kit (Bio-Rad; Carlsbad, CA). E. Original immunoblot probed with the anti-CaSeq1 antibody. F. Original immunoblot probed with the anti-GAPDH antibody.
